# CD4 T cells and neutrophils contribute to epithelial-mesenchymal transition in breast cancer

**DOI:** 10.1101/2023.02.15.528594

**Authors:** Amélien Sanlaville, Aurélien Voissière, Dominique Poujol, Margaux Hubert, Suzanne André, Clémence Perret, Jean-Philippe Foy, Nadège Goutagny, Marine Malfroy, Isabelle Durand, Marie Châlons-Cottavoz, Jenny Valladeau-Guilemond, Pierre Saintigny, Alain Puisieux, Christophe Caux, Marie-Cécile Michallet, Isabelle Puisieux, Nathalie Bendriss-Vermare

## Abstract

Epithelial-mesenchymal transition (EMT) is a central oncogenic mechanism, contributing both to transformation and metastatic dissemination. Inflammation and innate immune cells are known to favor EMT induction, but the role of adaptive immunity still remains unclear. Using an original murine mammary tumor model in immune cell subpopulation depletion experiments, we demonstrated that tumor cells maintain their epithelial phenotype in mice deficient for adaptive immune response, but undergo EMT in the presence of T-cells. This phenotypic conversion involves the major contribution of CD4 T cells, but not CD8 T cells nor B cells, undoubtedly demonstrating the pro-EMT role of CD4 T cells specifically among adaptive immune cells. Moreover, combined intra-tumor immune infiltrate and transcriptomic analyses of murine mammary tumors with various EMT phenotype revealed an inverse correlation between mesenchymal tumor cell and intratumoral neutrophil proportions, due to the reduced ability of mesenchymal cells to recruit neutrophils. Last, selective *in vivo* depletion of neutrophils and transcriptomic analysis of human breast tumor cohorts demonstrated the pro-EMT role of neutrophils and suggest a cooperation with CD4 T cells in EMT promotion. Collectively, our data highlight a novel mechanism of EMT regulation by both innate and adaptive immune compartments.

## INTRODUCTION

Epithelial-Mesenchymal Transition (EMT) is a fundamental transdifferentiation program enabling the reprogramming of polarized epithelial cells towards a mesenchymal phenotype (1,2). EMT plays crucial roles in embryonic development and physiological homeostatic response to tissue injury, but also in pathologies such as organ fibrosis and cancer progression (3,4). EMT is characterized by the loss of cell-to-cell contacts and a profound remodeling of the cytoskeleton, together resulting in the loss of apical-basal polarity and acquisition of fibroblast-like morphology. EMT is associated with a decrease in epithelial-specific gene expression, including E-cadherin, cytokeratins and EpCAM, and a gain in mesenchymal markers like vimentin, fibronectin and N-cadherin (5). EMT is orchestrated by a network of transcription factors (EMT-TFs), including members of the TWIST, ZEB and SNAIL families that act as genuine oncoproteins, fostering malignant transformation and primary tumor growth by alleviating key oncosuppressive mechanisms (6,7) and by providing cells with stemness features and resistance to chemotherapy and radiotherapy (8–10).

To better understand EMT process in the context of cancer, it may be important to consider that tumor development is not only limited to the tumor cell itself but also relies on its interactions with the microenvironment, and in particular with the immune system. Inflammation plays a key role at different stages of tumor development, including initiation, malignant conversion and metastasis (11,12). In particular, inflammatory cytokines such as TGF-β (13,14), TNF-α (15–17) or IL-6 (18–20), and innate immune cells including tumor-associated macrophages (TAM) (21,22) or myeloid-derived suppressor cells (MDSC) (23–25), are able to activate different signaling pathways finally converging on EMT induction and promoting tumor development and metastatic disease. In addition, EMT could be also a mechanism for cancer cells to evade the immune surveillance through favoring of an immune suppressive microenvironment, allowing immunosuppressive dendritic cell (DC) generation, regulatory T cell (T_reg_) promotion and resistance to CD8 T-cell-mediated lysis (26,27). These features led to wonder about possible EMT induction in response to immune pressure. If pro-EMT role of innate immunity has been quite well described, the involvement of adaptive immune response in EMT promotion has been poorly examined.

To investigate the role of anti-tumor adaptive immune response in EMT induction of tumor cells, we first developed original epithelial mammary tumor cell lines with a stable and EMT-independent expression of *Her2/Neu* oncogene and able to commit into EMT process *in vitro* upon TGF-β1 stimulation. *In vivo*, the growth of tumor cells was severely slowed when implanted in wild-type (WT) immunocompetent mice, as compared to immunodeficient RAG1KO and in CD4- and CD8-double-depleted mice. This delay observed in WT mice paralleled the acquisition of a mesenchymal phenotype while tumor cells sustained epithelial characteristics in immunodeficient mice. Interestingly, CD4 T cell but not CD8 T cell depletion restores tumor cell epithelial phenotype, demonstrating that CD4 T cells mediate an adaptive immune response leading to EMT induction. Intriguingly, neutrophil infiltrate was strongly reduced in mesenchymal tumors compared to epithelial ones, due to loss of neutrophil recruitment capacity in tumor cells undergoing EMT. Neutrophil depletion was also able to rescue epithelial phenotype of tumor cells, suggesting that both CD4 T cells and neutrophils favor EMT in breast cancer.

## RESULTS

### Establishment of a relevant murine mammary tumor cell line stably expressing Her2 and able to undergo EMT *in vitro*

It was previously shown that adaptive immune pressure against an overexpressed tumor-associated antigen (TAA) results in immunoediting through EMT induction (28–30). However the model used in these studies was based on the overexpression of a TAA under the control of mouse mammary tumor virus (MMTV) promoter that has an epithelial specific tropism (31), leading to TAA extinction during the EMT process and thus representing a major bias of these studies. NEU8.2 and NEU15 (32) tumor cell lines, previously established from spontaneous mammary tumors arisen in MMTV-*Her2/Neu* mice (**Supplementary Fig. 1a**), displayed typical cobblestone-like epithelial cell morphology (**Fig. 1a** and **Supplementary Fig. 1b**). TGF-β1 treatment induced striking morphological changes in NEU8.2 that acquired a mesenchymal-like spindle-shaped morphology and concomitantly lost Her2 expression, whereas NEU15 kept epithelial morphology as well as Her2 expression (**Fig. 1a** and **Supplementary Fig. 1b**). These results are in agreement with an epithelial cell-restrained MMTV-dependent Her2 expression and its extinction in mesenchymal cells, representing a major limitation of previous studies (28–30).

**Fig. 1:**
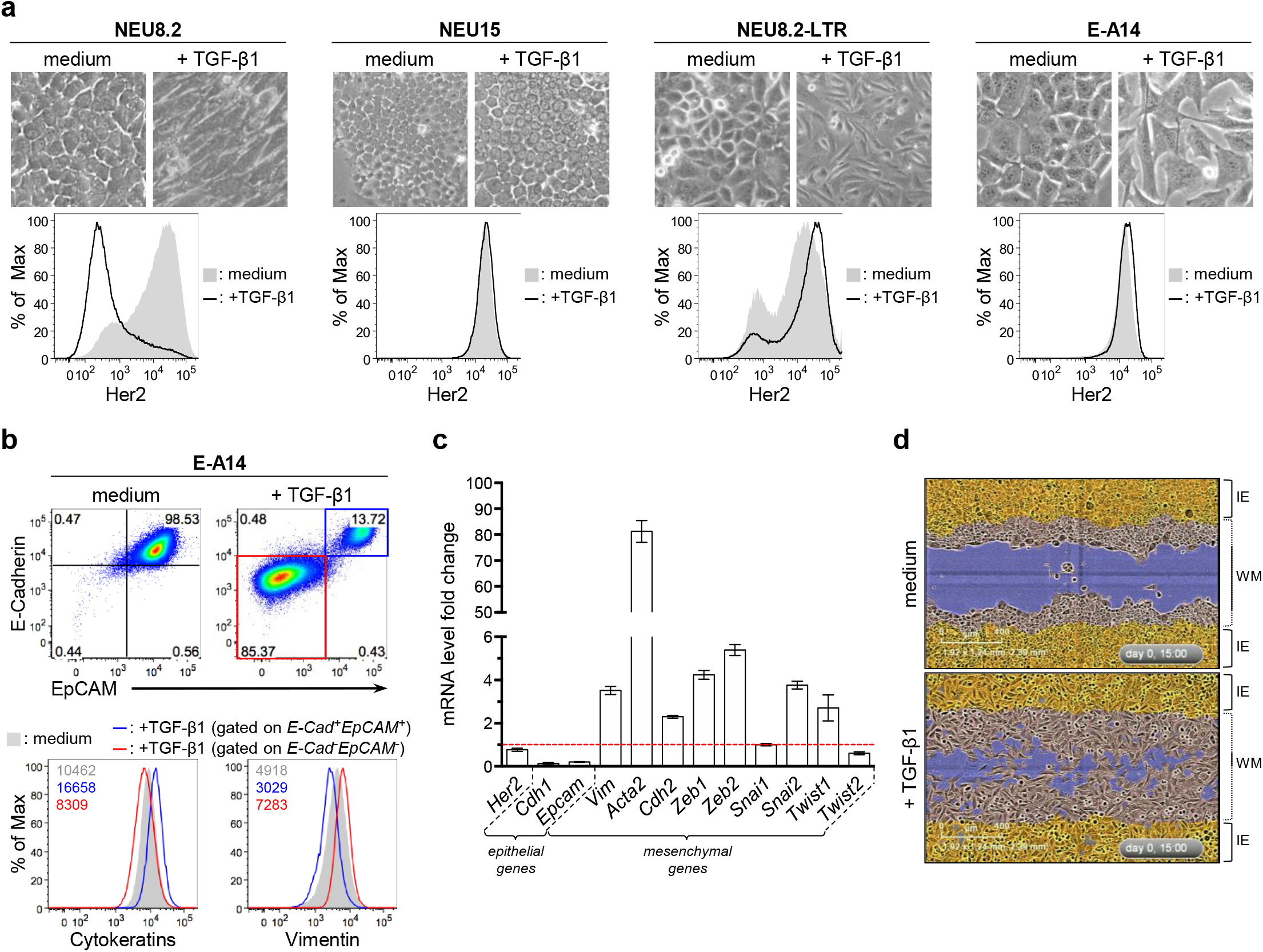
E-A14 is a plastic mammary tumor cell line that undergoes EMT in response to TGF-β1 treatment *in vitro*. **a-** NEU8.2, NEU15, NEU8.2-LTR and E-A14 mammary tumor cell lines were treated *in vitro* with TGF-β1. The morphology of cells was visualized using light microscopy (original magnification x200). Cells were also analyzed by flow cytometry for the expression of Her2. **b-** E-A14 cells were also analyzed by flow cytometry for the expression of E-Cadherin, EpCAM, Cytokeratins and Vimentin after TGF-β1 treatment. Gates are based on the isotype control corresponding to each marker. Numbers on histograms indicate mean fluorescence intensity. **c-** *Her2, Cdh1, Epcam, Vim, Acta2, Cdh2, Zeb1, Zeb2, Snai1, Snai2, Twist1* and *Twist2* mRNA expression was quantified by RT-qPCR in E-A14 cells after TGF-β1 treatment. Graph shows fold change of gene expression analyzed in TGF-β1-treated cells compared to control (medium) cells after normalization of gene expression by *Hprt1* and *Rplp0* housekeeping genes. Bars and error bars represent the mean and standard deviation, respectively. n=4. Red dotted line indicates a fold change of 1. **d-** E-A14 cells were grown in a monolayer in TGF-β1-supplemented medium. A scratch wound was made and images were taken using the Incucyte ZOOM instrument (Essen BioScience). Pictures presented here display wound healing after 15h. The blue region denotes the scratch wound mask (WM) over time as cells migrate into the wound region. The initial migrating/invading edges (IE) of the wound, created immediately following wound generation, are shown in yellow. TGF-β1: transforming-growth factor beta 1.

To bypass this promoter-dependent TAA expression issue that prevents from a proper evaluation of EMT-related interactions between adaptive immune cells and tumor cells, NEU8.2 was stably transduced with same *Her2* oncogene, but this time under the control of the ubiquitous Moloney murine leukemia virus (MoMuLV) LTR promoter. This allows to generate an epithelial cell line, named NEU8.2-LTR, retaining Her2 expression even after EMT induction by TGF-β1 (**Supplementary Fig. 1a** and **Fig. 1a**). An homogenous tumor clone, named E-A14 and obtained by limiting dilution, displayed an epithelial phenotype, characterized by cobblestone-like morphology, high expression of E-Cadherin/*Cdh1*, EpCAM/*Epcam* and cytokeratin molecules, and a very weak expression of the mesenchymal markers *Vim, Acta2, Cdh2* and EMT-TFs from *Zeb, Twist* and *Snail* families (**Fig. 1b-c** and **Supplementary Fig. 1a, d-e**). These epithelial features were comparable to the completely epithelial cell line NEU15, resistant to EMT induction by TGF-β1 (**Supplementary Fig. 1b, e**). Interestingly, TGF-β1 treatment induced EMT-like changes in E-A14 cell morphology towards a mesenchymal-like spindle-cell shape and increased inter-cellular separation, downregulation of E-Cadherin and EpCAM mRNA and proteins and over-expression of EMT markers (*Vim, Acta, Cdh2)* and EMT-TF (*Zeb1, Zeb2, Snai2, Twist1*) mRNA (fold change between 2.3 to 81.3; **Fig. 1a-c** and **Supplementary Fig. 1d-e**). This TGF-β1-induced mesenchymal phenotype resembles to that of 2G2, a clone isolated from another MoMuLV-*Her2*-transduced cell line (**Supplementary Fig. 1a**) and displaying a complete mesenchymal phenotype regardless of TGF-β1 treatment (**Fig. 1a-c** and **Supplementary Fig. 1c-e**). Moreover, a small proportion of E-A14 tumor cells kept E-Cadherin and EpCAM expression after TGF-β1 treatment, and coherently E-Cadherin^−^EpCAM^−^ E-A14 cells had lower expression of cytokeratins and increases expression of vimentin compared to E-Cadherin^+^EpCAM^+^ E-A14 cells consistent with EMT induction (**Fig. 1b**). In scratch-wound assay, NEU15 cells did not display invasive capacity regardless of TGF-β1 treatment and were only able to fill the wound by proliferation from the two edges, whereas 2G2 cells intensively migrated in the wound (**Supplementary Fig. 1f** and **Supplementary Files 1-4**), confirming their opposite epithelial and mesenchymal phenotype, respectively. Confirming its ability to commit into EMT, E-A14 non-invasive cells acquired mobility through TGF-β1-treatment (**Fig. 1d** and **Supplementary Files 5-6**). Importantly, Her2 was highly expressed and conserved in all the MoMulLV-*Her2*-transduced cell lines (E-A14 and 2G2), validating the EMT-independent Her2 expression in these newly generated cellular tumor models (**Fig. 1a, c** and **Supplementary Fig. 1c-d**). Altogether, these results indicate that E-A14 is a plastic Her2^+^ epithelial tumor cell line, able to undergo EMT *in vitro* upon TGF-β1 treatment and with a stable expression of the driving *Her2* oncogene.

### Adaptive immune response favors EMT *in vivo*

After demonstrating the relevance of E-A14 as a plastic EMT model, E-A14 was used to study EMT *in vivo*. E-A14 tumor growth was tightly controlled in WT mice (tumor volume: mean ± sd = 583mm^3^ ± 218 at day 20 post-injection), with a 30-60 days-long phase of equilibrium at the end of which tumors were rejected in almost half of mice whereas others escaped and progressed with a brisk growth curve. In contrast, E-A14 rapidly form tumors in all T and B cell-deficient RAG1KO mice reaching ethical end-point volume 20 days after tumor cell injection (2664mm^3^ ± 316) (**Fig. 2a**). This was also the case when tumor cells were injected in transgenic MMTV-*Her2/Neu* mice, with a tolerant immune system against xenogeneic rat Her2 (data not shown), suggesting that tumor rejection in WT mice is mainly due to anti-Her2 immune response. Taken together, these data demonstrate the ability of adaptive immune response to induce tumor rejection or tumor control until tumors escape.

**Fig. 2:**
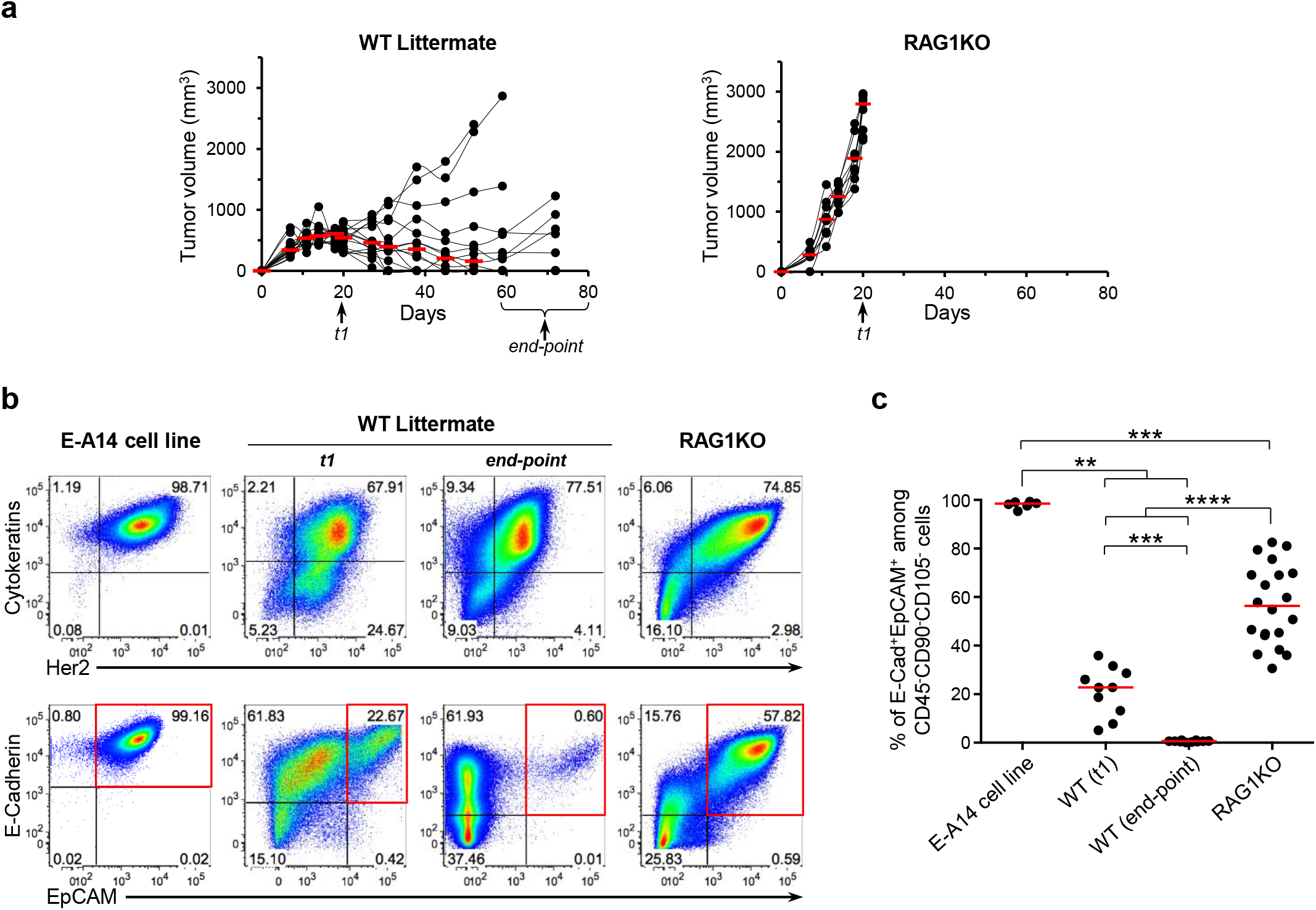
EMT process relies on adaptive immune response. **a-** E-A14 epithelial mammary tumor cells were injected into the mammary fat pad of RAG1KO (n=15) or WT littermate (n=9) mice. Tumor diameters were measured. Red bars indicate median tumor volume of the group. Arrows indicate sampling times at which some of the tumors were harvested (t1 = 18-21 days post-injection for RAG1KO and WT mice; end-point = end-point of tumor growth in WT mice, *i*.*e*. tumor volume >2500mm^3^) and **b-c-** analyzed by flow cytometry for Cytokeratins, Her2/Neu, E-Cadherin and EpCAM expression. Here are presented **b-** representative dot plot analysis and **c-** associated cumulative data graph after exclusion of CD45^+^ immune infiltrate, CD90^+^ fibroblasts and CD105^+^ endothelial cells. In comparison, expression of same markers was analyzed on *in vitro* cultured E-A14 tumor cell line at the injection time. **b-** Gates are based on the isotype control corresponding to each marker. **c-** Red bars on the graph indicate the median for each group. E-A14 cell samples (n=6), WT littermate mice harvested at t1 (n=10) and at end-point (n=10), and RAG1KO mice (n=20) were examined over at least three independent experiments. Kruskal-Wallis test. **, ***, ****: p ≤ 0.01, 0.001 or 0.0001. WT: wild-type.

Tumors were harvested 20 days after injection in each group (*t1*) (end time point for RAG1KO group), and also at the end point (after the immune evasion phase) in WT hosts (**Fig. 2a**). Importantly, Her2 expression was maintained in tumor cells during the tumor growth regardless of the host immune context, validating the relevance of this tumor cell model in the study of EMT *in vivo* (**Fig. 2b**). Moreover, the perfect co-expression between Her2 and cytokeratin molecules confirms the accuracy of the gating strategy using CD45^+^CD90^+^CD105^+^ cell exclusion to correctly identify tumor cells (**Fig. 2b**). Intriguingly, E-A14 tumors, obtained after injection of a completely epithelial E-Cadherin^+^EpCAM^+^ (E) cell population, displayed a mixt epithelial and mesenchymal phenotype 20 days after injection. While tumor cells mainly remain epithelial (56.9% ± 3.7 E-Cadherin^+^EpCAM^+^) in RAG1KO at day 20, this epithelial contingent represented only 21.2% ± 3.2 in WT hosts at the same time point and disappeared almost totally at the later end-point (0.6% ± 0.1) to be replaced by mesenchymal tumor cells (**Fig. 2b-c**). Therefore, these results demonstrate that, in WT mice, T and/or B cell-mediated adaptive immune pressure promotes EMT. Confirming these observations, two tumor cell lines (named M-A14.1 and M-A14.2) derived *in vitro* from E-A14 tumors harvested in WT hosts (**Supplementary Fig. 1a**), displayed a typical mesenchymal phenotype and a fibroblast-like morphology (downregulation of epithelial markers and strong overexpression of mesenchymal vimentin and EMT-TF, compared to E-A14), similar to that of 2G2 mesenchymal clone (**Supplementary Fig. 2**). Taken together, all these results demonstrate that adaptive immune response mediated by T and/or B cells is involved in EMT promotion.

### T cell-mediated immunity promotes EMT *in vivo*

In order to identify which immune cell subpopulation drives the mesenchymal conversion of tumor cells *in vivo*, E-A14 cell line was injected into B cell-(CD20 depletion) or T cell-(CD4 and CD8 double depletion) depleted mice compared to the corresponding control antibody depleted mice. As observed in RAG1KO mice (**Fig. 2a**), E-A14 tumor growth was highly accelerated in CD4 and CD8 double depleted mice compared to control mice (2383mm^3^ ± 626 versus 417mm^3^ ± 216, respectively, at day 20 post-injection). Conversely, CD20-mediated B cell depletion that completely abolished anti-tumor humoral response like CD4 and CD8 double depletion (**Supplementary Fig. 3**), did not modify E-A14 tumor growth (**Fig. 3a-b**) (1547mm^3^ ± 750 for CD20-depleted mice *versus* 897mm^3^ ± 643 for control mice at day 27 post-injection), suggesting that mainly T cell-mediated but not B cell-mediated immune response is responsible of tumor control and selection. While CD20 depletion did not prevent the acquisition of mesenchymal features by E-A14 tumor cells observed in immunocompetent control mice (**Fig. 3c-d**), CD4 and CD8 double depletion restored the epithelial tumor cell contingent (34.6% ± 3.9 in CD4/CD8-depleted mice *versus* 11.8% ± 4.2 at t1 and 0.2% ± 0.1 at the end-point in control mice) (**Fig. 3e-f**), demonstrating that T cell-mediated immunity favors EMT *in vivo* without any involvement of B cells.

**Fig. 3:**
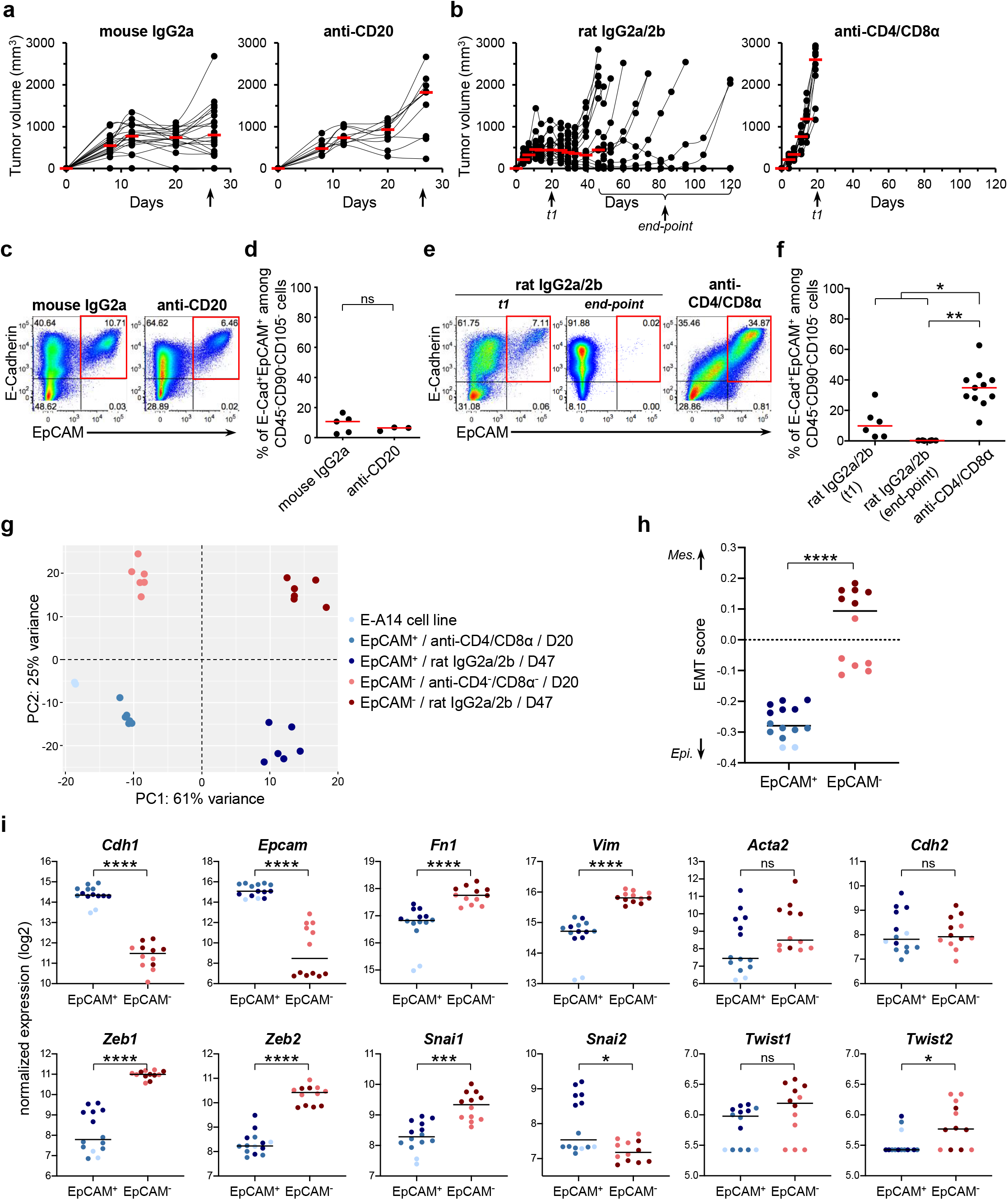
Adaptive T cell-mediated immunity is required for EMT induction of tumor cells *in vivo*. **a-b-** E-A14 epithelial mammary tumor cells were injected into the mammary fat pad of **a-** CD20-depleted (n=10) or isotype (mouse IgG2a) control-injected (n=20) mice, and **b-** CD4/CD8α double-depleted (n=10) or isotype (rat IgG2a and IgG2b) control-injected (n=19) mice. Tumor diameters were measured. Red bars indicate median tumor volume of the group. Arrows indicate sampling times at which some of the tumors were harvested (**c-d-** 29 days post-injection for anti-CD20 and corresponding control mice; **e-f-** t1 = 20-27 days post-injection; end-point = end-point of tumor growth in control mice, *i*.*e*. tumor volume >2500mm^3^) and **b-c-** analyzed by flow cytometry for E-Cadherin and EpCAM expression. Here are presented **c, e-** representative dot plot analyses and **d, f-** associated cumulative data graphs after exclusion of CD45^+^ immune infiltrate, CD90^+^ fibroblasts and CD105^+^ endothelial cells. Gates are based on the isotype control corresponding to each marker. Red bars on the graph indicate the median for each group. **c-d-** CD20-depleted mice (n=3) and control mice (n=5) were examined over one experiment. **e-f-** CD4/CD8 double-depleted mice (n=11) and control mice, harvested at t1 (n=6) and at end-point (n=6), were examined over at least two independent experiments. **d-** Mann-Whitney test and **f-** Kruskal-Wallis test. *, **: p ≤ 0.05 or 0.01. **g-i-** EpCAM^+^ and EpCAM^−^ tumor cells were sorted from E-A14 tumors harvested from CD4/CD8α double-depleted or isotype control-injected mice (n=6 each), when tumors reached their endpoint volume (at day 20 and day 47 post tumor injection, respectively). Transcriptomic analysis by RNA sequencing was performed on sorted cells and on 2 replicates of *in vitro* cultured E-A14 cell line. **g-** Principal-component analysis (PCA) plot considering the top 500 most variant genes. **h-** EMT score (signature defined by *Tan et al*., *EMBO Molecular Medicine 2014*) estimation by single sample gene set enrichment analysis (ssGSEA) for each cell sample. Black bars on the graph indicate the median for each group. Mann-Whitney test. ****: p ≤ 0.0001. **i-** Normalized expression of epithelial and mesenchymal genes compared between EpCAM^+^ and EpCAM^−^ isolated tumor cell samples. Black bars on the graph indicate the median for each group. Mann-Whitney test. *, ***, ****: p ≤ 0.05, 0.001 or 0.0001. EMT: epithelial-mesenchymal transition; Mes.: mesenchymal; Epi.: epithelial.

EPCAM^+^ and EpCAM^−^ tumor cells were then sorted from E-A14 tumors harvested at their respective growth end-point in CD4/CD8-depleted (20 days post-injection) and in immunocompetent control mice (47 days post-injection) to perform RNA sequencing analysis. Principal-component analysis perfectly segregates tumor samples according to the immune status of their host of origin (PC1) and their EMT phenotype (PC2) (**Fig. 3g**). Based on the transcriptomic data, an EMT score was then calculated for each tumor sample and also for *in vitro* cultured E-A14 cells. Independently of the host type from which tumor cells were harvested, EpCAM^−^ cells displayed a significantly higher mesenchymal-like EMT score (p<0.0001) as compared to EpCAM^+^ cells (**Fig. 3h**). Consistent with these data, the expression of *Cdh1* and *Epcam* epithelial genes was significantly reduced in EpCAM^−^ cells compared to EpCAM^+^ cells, while mesenchymal *Fn1, Vim, Zeb1/2, Snai1/2* and *Twist2* were highly enriched (**Fig. 3i**), validating that cell surface EpCAM loss, observed in tumors cells from immunocompetent mice, truly reflects EMT commitment. Altogether, these highly consistent results demonstrate that T cells favor EMT *in vivo*.

### CD4 T cell immune response induces EMT *in vivo*

Selective CD4 versus CD8 depletions were then performed. Compared to CD4/CD8-depleted mice, tumor growth was slightly decreased in CD4-depleted mice and CD8-depleted mice, and still strongly accelerated and homogenous among replicates compared to WT mice (**Fig. 4a**). Analysis of the tumor phenotype confirmed the acquisition by E-A14 epithelial tumor cells of a mesenchymal phenotype in WT hosts (12.2% ± 2.0 of E cells at t1 and 2.0% ± 0.6 at the end-point), and even in CD8 T cell-depleted mice (2.0% ± 0.6) (**Fig. 4b-c**). Interestingly, CD4 T cell depletion strongly impaired the mesenchymal conversion of tumor cells (47.2% ± 4.6 of E cells), even more than CD4/CD8 T cell-depletion (31.4% ± 2.3), demonstrating that CD4, but not CD8, T cell immune response favors tumor EMT *in vivo*.

**Fig. 4:**
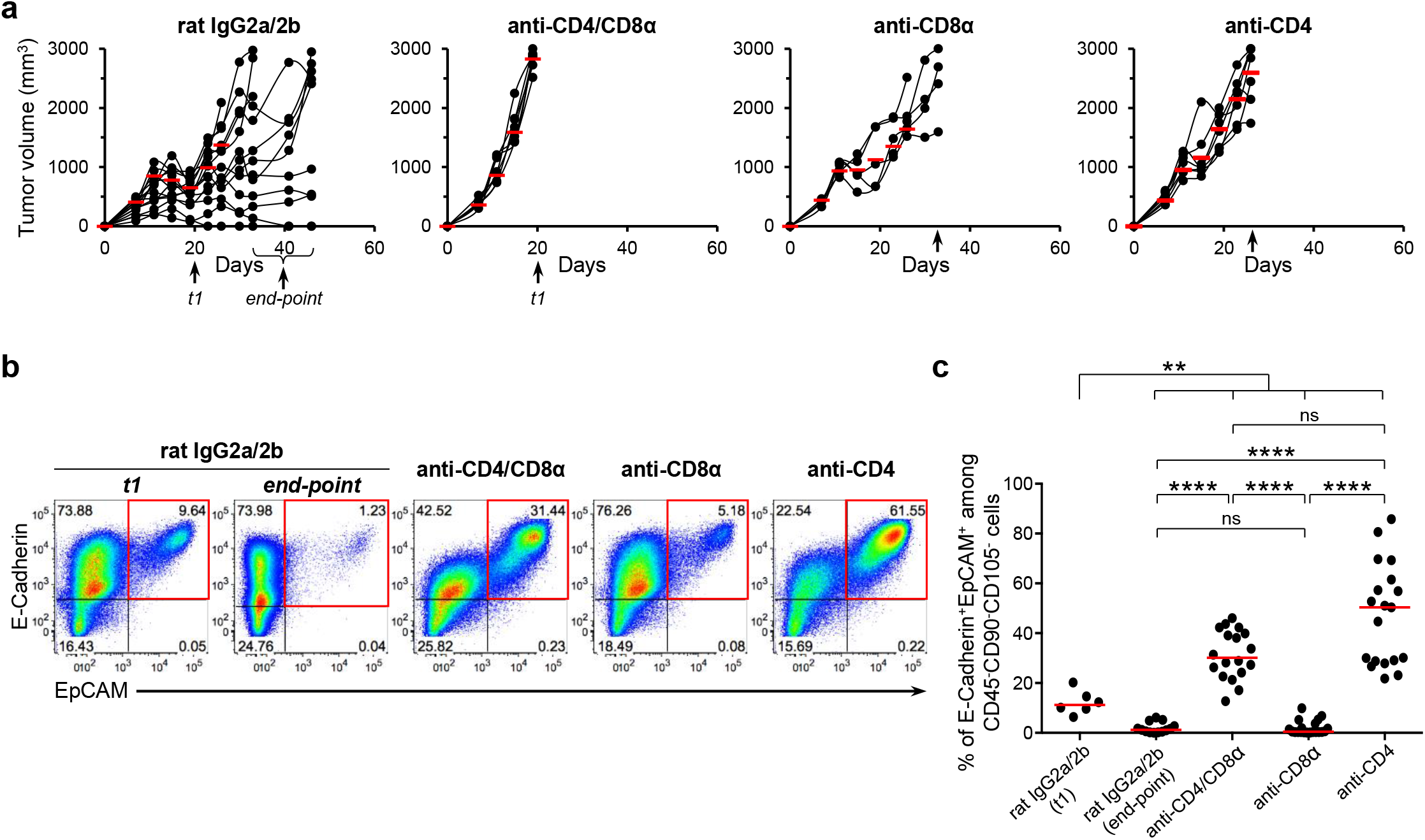
CD4 T cells are required for EMT *in vivo*. **a-** E-A14 epithelial mammary tumor cells were injected into the mammary fat pad of CD4/CD8α double-depleted (n=7), CD8α-depleted (n=5), CD4-depleted (n=7) or isotype (rat IgG2a and IgG2b) control-injected (n=14) mice. Tumor diameters were measured. Red bars indicate median tumor volume of the group. Arrows indicate sampling times at which some of the tumors were harvested (t1 = 20-22 days post-injection for CD4/CD8α double-depleted and control mice; 27-38 days post-injection for CD8α-depleted and CD4-depleted mice; end-point = end-point of tumor growth in control mice, *i*.*e*. tumor volume >2500mm^3^) and **b-c-** analyzed by flow cytometry for E-Cadherin and EpCAM expression. **b-** Representative dot plot analyses and **c-** associated cumulative data graph after exclusion of CD45^+^ immune infiltrate, CD90^+^ fibroblasts and CD105^+^ endothelial cells are presented. **b-** Gates are based on the isotype control corresponding to each marker. **c-** Red bars on the graph indicate the median for each group. Control mice harvested at t1 (n=6) and at end-point (n=13), CD4/CD8α double-depleted mice (n=18), CD8α-depleted mice (n=20) and CD4-depleted mice (n=19) were examined over at least two independent experiments. Kruskal-Wallis test. **, ****: p ≤ 0.01 or 0.0001.

### EMT modulates intra-tumor immune infiltrate by reducing neutrophil recruitment

E-A14 tumor immune infiltrate was then analyzed by flow cytometry. CD11b^+^ myeloid cells represent the dominant population regardless of the host immune context (**Fig. 5a**). Intriguingly, neutrophils frequency (defined as CD45^+^CD3^−^CD19^−^NKp46^−^CD11b^+^I-A/I-E^low/−^ Ly6G^+^Ly6C^+^ cells, not included in the other CD11b^+^ myeloid cell compartment) was higher in RAG1KO (16.0% ± 5.1 among CD45^+^ cells), double CD4/CD8-depleted (14.3% ± 4.1) and single CD4-depleted mice (16.9% ± 9.1), compared to control (7.1% ± 5.1) and CD8-depleted mice (5.9% ± 3.1) (**Fig. 5a-b**). Consistent with these data, single sample gene set enrichment analysis (ssGSEA) of human breast tumor samples, from The Cancer Genome Atlas (TCGA) (33) and METABRIC (34) cohorts, revealed that neutrophil enrichment score was significantly lower as CD4 T cell score enrichment increased (**Fig. 5c**), suggesting that neutrophils were preferentially enriched in tumors poorly infiltrated by CD4 T cells. Considering the phenotype of murine tumors with increased neutrophil frequency, neutrophils were preferentially enriched in epithelial polarized tumors (harvested in RAG1KO, double CD4/CD8-depleted and single CD4-depleted mice) compared to mesenchymal tumors (harvested in CD8-depleted or control mice) (**Fig. 5d**). These results correlate with pathway enrichment analysis of differentially expressed genes (DEGs) between mesenchymal EpCAM^−^ and epithelial EpCAM^+^ tumor cells, isolated from immunocompetent mice at tumor end-point as in **Fig. 3g-i**, and showing the highly significant downregulation of several gene pathways related to neutrophil or granulocyte chemotaxis and migration in mesenchymal cells (**Fig. 5e** and **Supplementary Fig. 4a**), all these gene pathways strongly correlating with neutrophil gene signatures (**Supplementary Fig. 4b**). This data suggests that EMT process induces tumor cell reprogrammation leading to the decrease of their neutrophil recruitment ability within the microenvironment. Thus, the effect of different tumor cell line supernatants on immune cell chemotaxis was then assessed using transwell migration assay. Interestingly, among all immune cells, neutrophils were the only immune cell subtype attracted by supernatants harvested from epithelial E-A14 and E-A21 (another clone originating from NEU8.2-LTR cell line) tumor cells, whereas supernatants from mesenchymal 2G2 and M-A14 cell lines did not induce any immune cell recruitment (**Fig. 5f** and **Supplementary Fig. 4c**). In line with this observation, the concentration of CXCL-1, one of the major attractants of neutrophils in the mouse (35–37), was detected at much higher levels in the supernatant of epithelial tumor cells compared to that of mesenchymal tumor cells (**Fig. 5g**), suggesting a decrease of CXCL-1 chemokine production during EMT process. Taken together, these results confirmed the down regulation of the neutrophil recruitment ability by tumor cells during EMT process.

**Fig. 5:**
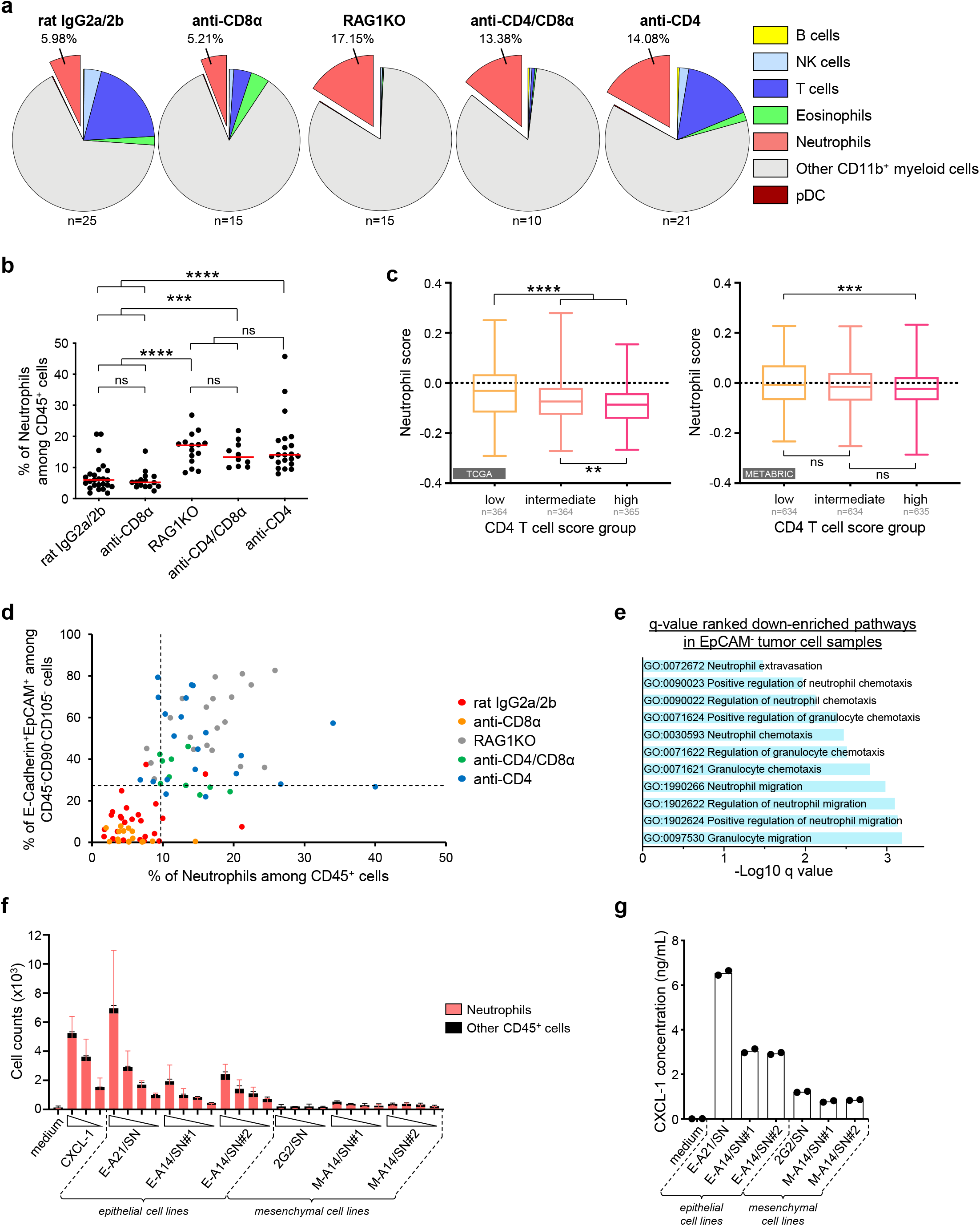
Neutrophil infiltration is strongly reduced in mesenchymal tumors due to loss of neutrophil recruitment ability by EMT tumor cells. **a-** Circular diagram of immune infiltrate subpopulations analyzed in tumors harvested after injection of E-A14 tumor cells into CD4/CD8α double-depleted (n=10), CD4-depleted (n=21) or CD8α-depleted (n=15), isotype (rat IgG2a and IgG2b) control-injected (n=25) and RAG1KO (n=15) mice. CD45^+^ immune cells were subdivided into B cells (CD19^+^), NK cells (NKp46^+^), T cells (CD3^+^), eosinophils (CD3^−^CD19^−^NKp46^−^CD11b^+^I-A/I-E^low/−^Ly6G^−^SiglecF^+^), neutrophils (CD3^−^ CD19^−^NKp46^−^CD11b^+^I-A/I-E^low/−^Ly6G^+^Ly6C^+^), other myeloid cells (CD3^−^CD19^−^NKp46^−^CD11b^+^), and plasmacytoid dendritic cells (CD3^−^CD19^−^NKp46^−^CD11c^+^SiglecH^+^). Numbers on diagrams indicate median frequency of neutrophils among CD45^+^ infiltrating immune cells for each group. **b-** Frequency of tumor-infiltrating neutrophils in E-A14 tumors harvested in CD4/CD8α double-depleted mice (n=10), CD4-depleted mice (n=21), CD8α-depleted mice (n=16), control mice (n=26) and RAG1KO mice (n=20) and determined by flow cytometry. Red bars on the graph indicate the median for each group. Kruskal-Wallis test. ***, ****: p ≤ 0.001 or 0.0001. c-Single sample enrichment scores of neutrophil signature (The human Protein Atlas) in transcriptomic data of breast tumors from human sample cohorts. Groups of breast tumors, from TCGA (left panel, n=1093 samples) and METABRIC (right panel, n=1903 samples), were defined on the basis of the memory CD4 T cell score of each sample determined by ssGSEA, allowing to define 3 groups corresponding to tertiles with low, intermediate and high memory CD4 T cell enrichment score. Kruskal-Wallis test. **, ***, ****: p ≤ 0.01, 0.001 or 0.0001. **d-** Percentage of epithelial (E-cadherin^+^EpCAM^+^) tumor cells versus frequency of tumor-infiltrating neutrophils in E-A14 tumors harvested in all the previous mouse models. Dotted lines correspond to the median values for epithelial tumor cell (27.3%) and tumor-infiltrating neutrophil (9.69%) frequencies of all samples. **a-b, d-** Data were obtained from at least three independent experiments. **e-** Pathway enrichment analysis of differentially expressed genes (DEGs) between EpCAM^−^ and EpCAM^+^ tumor cells, isolated from E-A14 tumors harvested in isotype (rat IgG2a/2b) control-injected mice (n=6) at tumor end-point (47 days after tumor cell injection). Here are presented down-enriched pathways related to neutrophils or granulocytes in EpCAM^−^ tumor cell samples (compared to EpCAM^+^). **f-** Dose-response effects of epithelial or mesenchymal tumor cell line supernatants on immune cell chemotaxis. Absolute counts of migrating neutrophils (CD45^+^Ly6G^+^Ly6C^int^) and other CD45^+^ immune cells are presented. For E-A14 or M-A14, two different supernatants (SN#1 and SN#2) were produced during distinct cell culture experiments. CXCL-1 was used as a positive control. Bars and error bars represent the mean and 95% confidence interval, n=3. Data are representative of two independent experiments. **g-** Protein concentration of CXCL-1 were determined by multiplex assay in the cell culture supernatants used in **f-**. Bars represent the mean.

### Neutrophils and CD4 T cells cooperate to favor EMT *in vivo*

To fully characterize the link between neutrophils and epithelial/mesenchymal phenotype of tumor cells, E-A14 tumor cells were implanted into anti-Ly6G-neutrophil-depleted mice. Interestingly, tumors harvested in anti-Ly6G mice displayed a more epithelial phenotype (43.1% ± 4.9 of E-Cadherin^+^EpCAM^+^ cells) compared to control mice (18.3% ± 3.6), and similar to that observed in CD4-depleted mice (57.2% ± 7.7), demonstrating that neutrophils are also responsible of EMT commitment *in vivo* (**Fig. 6**). Ly6G and CD4 co-depletion did not significantly increase the proportion of epithelial tumor cells (63.1% ± 6.6), suggesting a redundant or cooperative role of CD4 T cells and neutrophils in EMT promotion *in vivo* rather than an additive effect. Consistently, ssGSEA of human breast tumor samples, from METABRIC (**Fig. 6c**) cohort, revealed that EMT enrichment score (based on EMT signature defined by Tan *et al*. (38)) was significantly higher in tumors with the highest enrichment score of both neutrophils and CD4 T cells populations, lower in tumors with the lowest enrichment score of either immune cell subpopulation, all the more in tumors with the lowest enrichment score for both subpopulations. This was confirmed by the analysis of breast tumor samples from TCGA cohort and also using another EMT signature (EMT Hallmark) (**Supplementary Fig. 5**). Taken together, these results suggest an association between mesenchymal phenotype of the tumor and the presence of both tumor-infiltrating CD4 T cells and neutrophils, strongly supporting the notion that CD4 T cells and neutrophils cooperate to favor EMT *in vivo*.

**Fig. 6:**
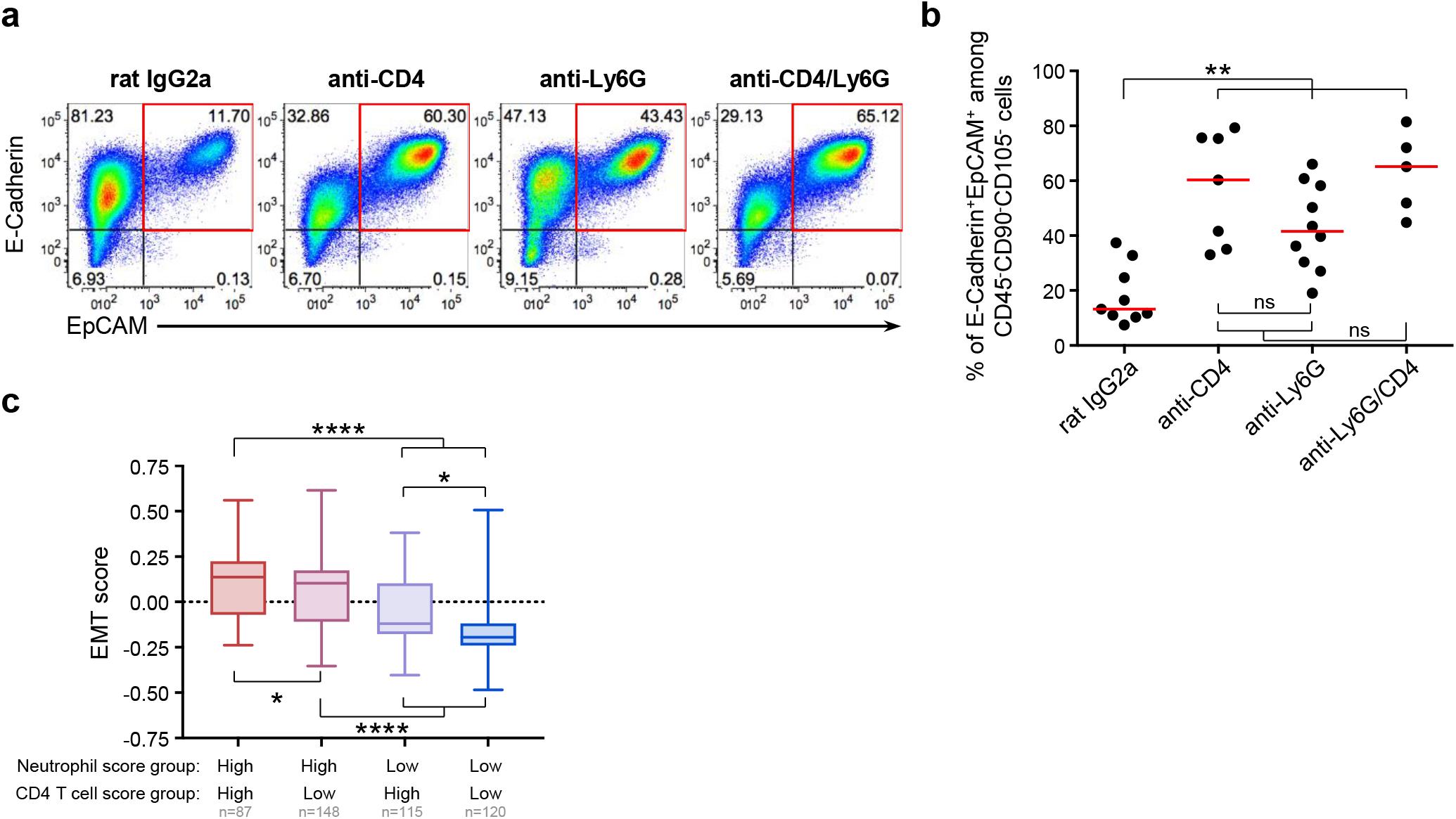
Neutrophils and CD4 T cells cooperate in EMT induction of tumor cells *in vivo*. E-A14 epithelial mammary tumor cells were injected into the mammary fat pad of isotype (rat IgG2a) control-injected, CD4-depleted, Ly6G-depleted or CD4/Ly6G double-depleted mice. Tumors were harvested at 20-22 days after tumor cell injection for each group and analyzed by flow cytometry for E-Cadherin and EpCAM expression. **a-** Representative dot plot analyses and **b-** associated cumulative data graph after exclusion of CD45^+^ immune infiltrate, CD90^+^ fibroblasts and CD105^+^ endothelial cells are presented. Gates are based on the isotype control corresponding to each marker. Red bars on the graph indicate the median for each group. Control mice (n=9), CD4-depleted mice (n=7), Ly6G-depleted mice (n=10) and CD4/Ly6G double-depleted mice (n=5) were examined over two independent experiments. Kruskal-Wallis test. **: p ≤ 0.01. **c-** Single sample enrichment scores of the EMT signature (defined by *Tan et al*., *EMBO Molecular Medicine 2014*) in transcriptomic data of breast tumors from METABRIC cohort. Tumors were selected according to their neutrophil and memory CD4 T cell enrichment scores (also determined by ssGSEA), with high and low groups defined by the 25% of tumors with the highest and the lowest scores. Kruskal-Wallis test. *, ****: p ≤ 0.05 or 0.0001.

## DISCUSSION

Here we demonstrated, using a relevant mouse model based on tumor cell lines plastic for EMT and stably expressing Her2 tumor-associated-onco-antigen, that adaptive immune system induces EMT in vivo. Using selective immune cell depletion, we excluded a role for B cells while highlighting the contribution of T cells, and specifically CD4 T cells without any CD8 T cell involvement, in this phenotypic transdifferentiation. Furthermore, immune cell infiltrate analysis in mouse mammary tumors revealed an association between a deprivation in intratumoral neutrophils and a mesenchymal tumor phenotype, explained by the downregulation of gene clusters involved in neutrophil recruitment in the course of EMT. These observations were complemented by the analysis of public human breast cohorts demonstrating a strong association between mesenchymal phenotype of the tumor, and CD4 T cells and neutrophils tumor infiltration. Finally, our study highlighted a previously unknown role of neutrophils in EMT induction, possibly through their cooperation with CD4 T cells.

Our results clarify the role of CD4 and CD8 T cell sub-populations in EMT induction *in vivo*, contrasting with previous studies alternatively showing that each one could be involved in EMT promotion (28–30). This discrepancy might result from the lability of tumor-driving Her2 oncogene expression during EMT in the MMTV-*Her2/Neu* mouse model used in these studies, MMTV promoter being switched off in mesenchymal cells.

The major original contribution of this work resides in the identification that the adaptive immune response favors EMT and that this deleterious impact relies on CD4 T cells but not on CD8 T cells or humoral response. Since CD4 T cells orchestrate, directly or indirectly, a wide range of immune functions, it is necessary to decipher their precise contribution. Among CD4 T cells, regulatory T cells (Treg) could be involved in EMT induction through their production of TGF-β (39). Using CD25 antibody to deplete Treg, we observed partial restoration of tumor epithelial phenotype (mean percentage of epithelial cells [95% confidence interval]: 27.57% [14.35; 40.78] in CD25-depleted mice; 40.45% [21.69; 59.21] in CD4-depleted mice; 12.20% [7.18; 17.22] in control IgG1-depleted mice), suggesting that Treg participate to EMT promotion *in vivo*. However, to avoid depletion of CD25-overexpressing activated effector T cells, CD25 depletion was performed just prior tumor injection, thus with a time-limited efficacy. The role of Treg in EMT promotion was also suggested in another study, demonstrating that Treg induce fibronectin overexpression on pulmonary epithelial cells *in vitro*, and showing that anti-CD25 depletion alleviated the mesenchymal phenotype observed in a post-irradiation fibrosis-induction pulmonary model (40). An accurate method to answer this question in a tumor context would be to use the FoxP3-DTR mouse model to specifically deplete Treg through tumor development and analyze the impact on tumor cell phenotype, however these mice are not available in a FVB genetic background.

Another CD4 T cell subpopulation potentially involved in EMT induction are Th17 lymphocytes via IL-17 secretion. The contribution of IL-17 in tumorigenesis is still unclear between pro and anti-tumor function. Several studies point to a role in EMT induction: i) an increased migration and invasion of breast cancer cells after IL-17α treatment (41), ii) a preferential Th17 cell infiltrate in triple-negative breast cancer, a subtype of tumors associated with EMT-features (41), iii) the IL-17-dependent acquisition of EMT hallmarks by pulmonary epithelial cells (42) and iv) the decrease of EMT features observed in lung tumor cells after IL-17 cytokine neutralization (43). Pro-EMT effect of IL-17 could be due to its ability to activate STAT3 (44) or NF-κB (45,46), two converging points of multiple cytokine-induced signaling pathways leading to the activation of EMT-TF (47–50). A role of Th17 cells would be consistent with the role of neutrophils as IL-17 is well known to trigger the production of neutrophils chemoattractants (51–55) and also to regulate neutrophil homeostasis through granulocyte colony-stimulating factor (G-CSF) (56). However, the fact that the depletion of CD4 T cells results in neutrophil accumulation argue against this hypothesis.

In our experiments, CD4 but not CD8 T cell depletion, was sufficient to maintain the epithelial tumor phenotype, but at most for a maximum of 50-60% of cells, meaning that EMT could be induced *in vivo* via other signals or partners. Several hypotheses could be formulated. First, tumor growth was tremendously fast in immunodeficient mice and associated with high cellular mortality notably in the center of the tumor, potentially leading to over-estimation of the percentage (among viable tumor cells) of mesenchymal tumor cells preferentially found in tumor periphery (57,58). Moreover, due to the rapid growth, vascularization could be not correctly established within the tumor, leading to hypoxic conditions, known to induce EMT (59–63). Another hypothesis involved innate immune cells that highly infiltrate tumors. Indeed, numerous studies showed that TAM or MDSC were able to induce EMT of tumor cells (21,24,64,65), probably through their secretion of multiple EMT-inducing cytokines such as TGF-β, TNF-α, IL-6 or IL-1 (14,15,18,19,66). TAM or MDSC could thus induce mesenchymal conversion of a non-negligible proportion of tumor cells even in the absence of T cells.

Interestingly, neutrophils also appear to be key players in EMT induction in our model. The immune infiltrate analysis of mouse mammary tumors revealed positive correlation between percentages of epithelial cells and neutrophils. This observation was confirmed by transcriptomic analysis of epithelial and mesenchymal tumor cells harvested *in vivo* and also by *in vitro* migration assays both emphasizing the reduced ability of mesenchymal cells to attract and recruit neutrophils. The neutrophil depletion experiments revealed that neutrophils plays a key role in EMT promotion *in vivo*, another major contribution of our work since the EMT-inducing role of neutrophils has been mentioned in few studies and only in vitro (67–69). The fact that the co-depletion of CD4 T cells and neutrophils leads to similar maintenance of epithelial tumor cell phenotype, compared to the single depletion of each of them, argues for a potential cooperation between neutrophils and CD4 T cells in EMT induction. This hypothesis is consistent with the maintenance of epithelial tumor phenotype while neutrophils accumulate in the tumor microenvironment in absence of CD4 T cells. The observed association of mesenchymal enrichment score with combined neutrophil and CD4 T cell scores in human breast tumors further strengthens this hypothesis. Altogether, these findings led us to propose a model whereby neutrophils are early attracted by epithelial tumor cells and cooperate with CD4 T cells to trigger EMT transdifferentiation process, thus resulting in the loss of chemoattractants for neutrophils by EMT cells and therefore lower neutrophil recruitment within the tumor (**Fig. 7a**). In another alternative model considering EMT as an immune evasion mechanism as described in some studies (26,27), the unique cooperation of CD4 T cells and neutrophils may be able to drive the efficient elimination of epithelial tumor cells only, leading to the selection and accumulation of previously transdifferentiated mesenchymal tumor cells (**Fig. 7b**). In these scenarios, CD4 T cell or neutrophil depletion leads to the accumulation of epithelial tumor cells since efficient immune cooperation is broken up, leading to epithelial tumor cell accumulation that are described to have proliferative advantages compared to mesenchymal tumor cells (1,70,71).

**Fig. 7:**
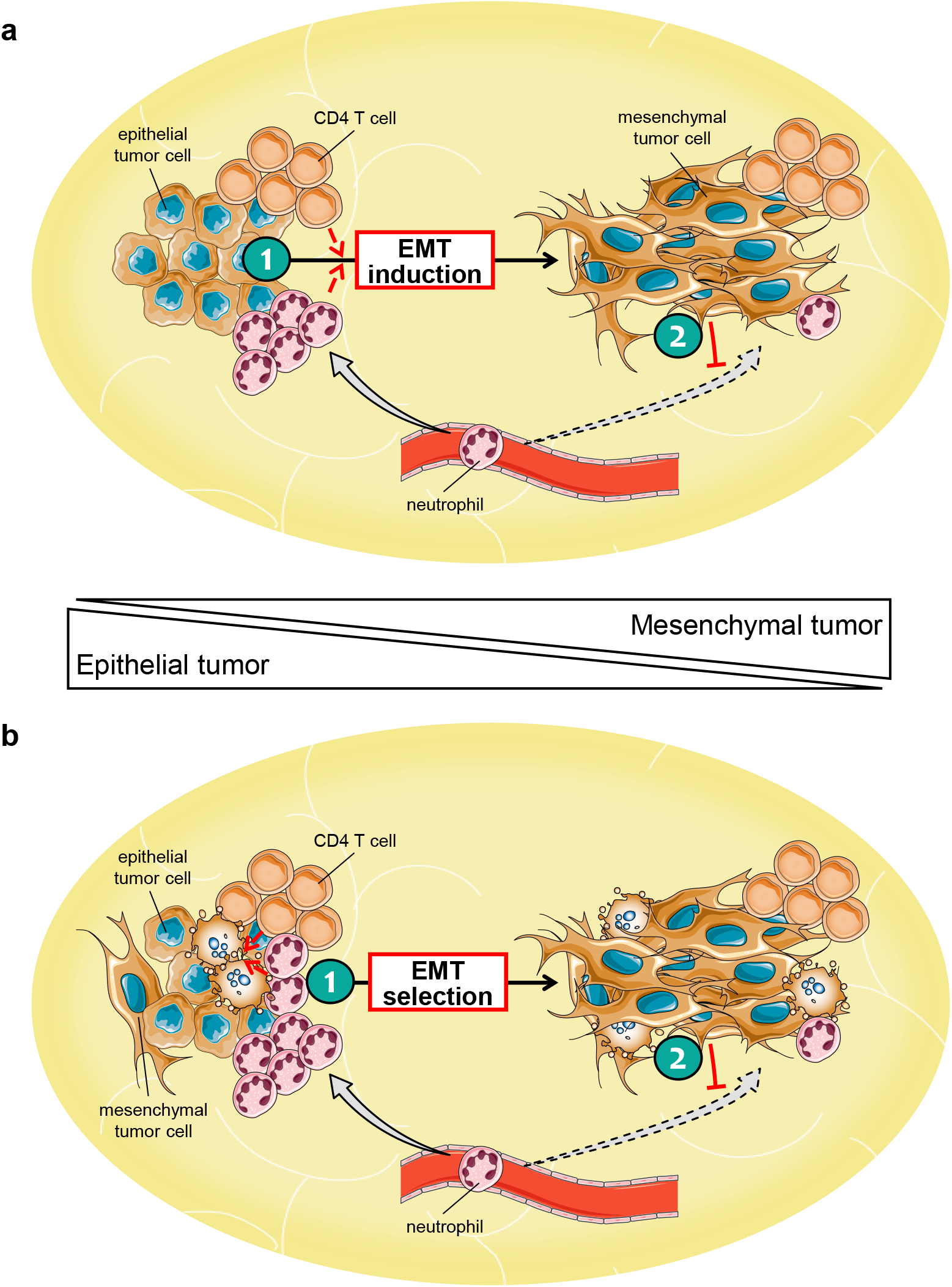
Cooperation between CD4 T cells and neutrophils in promotion of EMT in tumor cells. Two different models could be proposed to explain mesenchymal phenotype promotion by CD4 T cells and neutrophils *in vivo*. **a**-First, the concomitant presence of CD4 T cells and neutrophils (1) favors the induction of EMT transdifferenciation process in tumor cells (2), that in turn lose the ability to attract and recruit neutrophils. **b-** Alternatively, CD4 T cells and neutrophils (1) may preferentially target epithelial tumor cells, indirectly leading to (2) an enrichment of mesenchymal cancer cells within the tumor, preventing neutrophil recruitment. EMT: epithelial-mesenchymal transition.

All these observations are derived from analyses performed at a late tumor development stage. To complete these scenarios, it is also possible that at time of tumor cell injection, sentinel neutrophils may target epithelial tumor cells for lysis and phagocytosis (72–76). Cytokines secreted from anti-tumor neutrophils, massively recruited at this early time point, could be hijacked by the tumor cells and trigger their progressive EMT conversion, thus resulting in a concomitant reduction of anti-tumor neutrophil infiltration and generation of completely mesenchymal tumors devoid of neutrophil infiltration at a more advanced stage. Of note, triggering EMT as early as possible would give a selective advantage for tumor cells as EMT may be a potential powerful immune evasion mechanism. In this field, several studies have made their contribution showing that EMT endows tumor cells with immunosuppressive properties enabling dendritic cell alteration and regulatory T cell promotion (27), or resist to CD8 T cell lysis through alteration of the immune synapse formation and autophagy activation (26). Understanding the EMT-driven mechanisms leading to immune escape would help to design new therapeutic approaches to prevent metastatic development and relapse.

## METHODS

### Mice

Wild-type FVB/N (WT FVB) and FVB/N-Tg(MMTV*neu*)202Mul (77) female mice were purchased from Charles River Laboratory and The Jackson Laboratory respectively. RAG1KO mice in the FVB/N background were kindly provided by Dr K. de Visser (78) (Amsterdam, Netherlands). Animals were maintained in the specific pathogen free animal facility AniCan at the Centre Léon Bérard (Lyon, France). All animal procedures were reviewed and approved by the local Animal Ethic Committee (CECCAPP) and authorized by the French research ministry in accordance with the European directive 2010/63/UE.

### Tumor cell lines

NEU15 (32), NEU8.2 and NEU17 cell lines were established from short-time culture of spontaneously arising tumors in transgenic MMTV*neu* female mice (77) (**Supplementary Fig. 1a**). ANV17 cell line was previously obtained after orthotopic implantation in WT FVB mice of NEU17 cell line that led to the *in vivo* selection of a Her2 antigen-negative variant (ANV) that lost Her2 expression. All cell lines were grown in vitro at 37°C and 5% CO2 in DMEM (Life Technologies) supplemented with 10% heat inactivated fetal calf serum (FCS) (PAA), 200Units.mL^−1^ penicillin, 200µg.mL^−1^ streptomycin, 1% L-glutamine (all from Life Technologies), 2µg.mL^−1^ of puromycin (Invitrogen) for 2G2, E-A14 and M-A14 cells and 10µg.mL^−1^ of Insulin-transferrin-sodium selenite media supplement (Sigma-Aldrich) for NEU15.

### Generation of E-A14 and 2G2 cell lines

#### Production of pBabe LTR-Neu Retroviruses

The activated allele of rat *her2/neu* (kindly provided by Dr B.H. Nelson, Columbia, Canada) was amplified by PCR and inserted into the retroviral vector pBABE-puro (Addgene) containing the puromycin-resistance gene and the MoMuLV LTR promoter. This was followed by transformation into competent XLgold bacteria (Agilent) and amplification. The plasmid now referred as pBabe-LTR-Neu was further sequenced. pBabe retrovirus was produced by calcium phosphate transfection in Platinum-E cells (79) (Cell Biolabs), HEK 293 T cells stably expressing the retroviral GAG, POL and ENV components. After 72hrs of transfection, viral-particles-containing-supernatant was harvested and filtered through a 0.45µm cellulose filter. Finally, virus was concentrated using the 100000 MWCO Vivaspin® Concentrator (Sartorius Stedim) and titrated on mammary epithelial cells. Titer was 1.48.10^7^ infectious unit per milliliter.

#### Viral transduction

1.10^5^ of NEU8.2 or ANV17 cells/well were plated in 6-well dishes. After 24hrs, medium was replaced with freshly-made viral mixture (MOI=0.148) containing 0.8µL.mL^−1^ polybrene in 5% FCS complete medium. 24hrs later, viral medium was replaced with classical culture medium. Efficiently transduced cells, hereinafter referred as NEU-LTR cells, were selected by addition of 2µg.mL^−1^ puromycin before isolation by limiting-dilution cloning. E-A14 and 2G2 were thus obtained after cloning of NEU8.2-LTR and ANV17-LTR cells respectively.

#### In vitro EMT-induction assay

1.10^5^ cells/well were plated in 6-well dishes and treated with 10ng.mL^−1^ TGF-β1 (Peprotech) for 10 days. Then, cells were harvested, counted and stained with different antibodies (as previously described). Cell morphology was visualized with an Axiovert 25 inverted microscope (Zeiss).

#### Tumor cell line migration assay

NEU15, 2G2 and E-A14 cells were first treated or not with 10ng.mL^−1^ TGF-β1 for 10 days and then plated in a 96-well ImageLock plates (Essen Bioscience) at a density between 30 000 to 70 000 cells/well to obtain approximately 90% of confluence. 24h after plating, a scratch of about 1 mm was created using a 96-pin WoundMaker (Essen Bioscience, Michigan, USA) and the cell culture medium was replaced. Cells were still treated or not with TGF-β1. The scratch area was monitored every 30min using the Incucyte Live-cell Imaging System (Essen Bioscience) over a 35-50 hour time period using a 10x objective. The analysis was performed using the “wound confluence” from Incucyte ZOOM software, images and videos were exported using the same software. Each condition was performed in triplicates.

#### In vivo transplanted orthotopic mammary tumor models

6-8 week old mice were injected with 5.10^6^ tumor cells in 200µL PBS into the fourth mammary fat pad. Animals were checked twice a week for tumor growth. Tumor volume was calculated using the formula for an ellipsoid volume: V = π/6 x ab^2^. The two largest perpendicular tumor diameters ‘a’ and ‘b’ (a > b) were measured using a Vernier caliper. For ethical reason, the tumor-bearing mice were euthanized before the largest tumor diameter reached 17 mm.

#### Immune cell subset depletion experiments

For CD4 and/or CD8 T cell depletion experiments, 6-8 week old female mice were intraperitoneally (i.p.) injected twice per week with 100µg anti-CD4 (GK.1.5; BioXCell) and/or 100µg anti-CD8α (53-6.72; BioXCell) antibodies all along tumor growth. Treatments started 2 weeks before tumor implantation to ensure complete depletion. Purified rat IgG2a (2A3; BioXCell) and/or rat IgG2b (LTF2; BioXCell) were used as control.

For B cell depletion experiments, mice were i.p. injected once every two weeks with 500µg anti-CD20 (5D2; Genentech) antibody, starting 2 weeks and a half before tumor implantation and maintaining all along tumor growth. Purified mouse IgG2a (C1.18.4; BioXCell) was used as control.

For neutrophil depletion experiments, mice were i.p. injected twice per week with 100µg anti-Ly6G (1A8; BioXCell) antibody, starting 2 weeks before tumor implantation and maintaining all along tumor growth. CD4 T cell additional depletion was performed as described above. Purified rat IgG2a was used as control.

#### Detection of anti-tumor Ig in plasma of tumor-bearing mice

Plasma from E-A14 tumor-bearing mice were collected to determine anti-E-A14-specific immunoglobulins titers by standard indirect immunofluorescence. E-A14 cells grown *in vitro* were stained by serial dilutions of plasma for 15 min at 4°C. Cells were washed and incubated at 4°C with goat anti-mouse Ig-PE (SouthernBiotech). Cells were washed and mean fluorescence intensity (MFI) was measured by flow cytometry (FCM). The concentration of anti-E-A14-specific Ig in plasma was estimated in comparison with a standard curve generated with anti-Her2 monoclonal antibody (Ab-4; Calbiochem).

#### Ex vivo cell isolation

Tumors were collected at different time points and enzymatically digested in 10mg.mL^−1^ collagenase and 0.2mg.mL^−1^ DNAse I (Sigma-Aldrich) for 30min at 37°C. Red blood cells were then lysed 5min with Pharmlyse Buffer (BD Biosciences) at room temperature.

#### Antibodies, flow cytometry analyses and cell sorting

Multiparametric FCM analyses were performed on i) *in vitro*-cultured tumor cell lines, or on ii) single cell suspensions derived from harvested mammary tumors. The FCM panel used to analyze EMT phenotype of *in vitro* tumor cells relied on the use of the anti-mouse pan-cytokeratins (C-11; Biolegend), CD73 (TY/11.8; Biolegend), vimentin (D21H3; Cell Signaling Technology), E-cadherin (36/E-Cadherin; BD Biosciences), EpCAM (G8.8; Biolegend) antibodies. EMT phenotype of *in vivo* isolated tumor cells was analyzed using the same antibody panel implemented with anti-mouse CD45 (30-F11; Biolegend), CD90.1 (OX-7; Biolegend) and CD105 (MJ7/18; BD Biosciences) antibodies to exclude immune cells, fibroblasts and endothelial cells respectively from analysis. In both cases, Her2 expression was assessed using the custom-conjugated mouse anti-rat c-ErbB2/c-Neu (7.16.4; kind gift from Dr M. Greene; Philadelphia, Pennsylvania). Immune infiltrate of *in vivo* tumors was monitored using two different antibody panels. First was focused on lymphocyte subpopulation analysis and relied on the use of anti-mouse CD8β (H35-17.2; eBioscience), CD19 (6D5; Biolegend), CD4 (RM4-5; Biolegend), NKp46 (29A1.4; Biolegend), CD3ε (eBio500A2; eBioscience), CD45 (30-F11; Biolegend) and FoxP3 (FJK-16s; eBioscience), whereas the second allowed myeloid cell characterization with anti-mouse I-A/I-E (M5/114.15.2; Biolegend), CD11b (M1/70; Biolegend), Ly6G (1A8; BD Biosciences), SiglecH (551; Biolegend), CD3 (17A2; Biolegend), CD19 (6D5; Biolegend), NKp46 (29A1.4; Biolegend), SiglecF (E50-2440; BD Bioscience), CD45 (30-F11; Biolegend), CD11c (N418; Biolegend) and Ly6C (AL-21; BD Bioscience) antibodies. Each staining panel contained a viability marker to exclude dead cells from the analysis: LIVE/DEAD Fixable Dead Cell Stains (Life Technologies) or Zombie Fixable Viability Dyes (Biolegend). All stainings were performed in presence of Fc Block/anti-mouse CD16/32 (93; Biolegend).

All data were acquired with LSRFortessa or FACScan (BD Biosciences) cytometers and analyses were performed with FlowJo software (Tree Star) after gating on viable cells and doublet exclusion.

For mouse tumor cell transcriptomic analysis, Her2^+^EpCAM^+^ and Her2^+^EpCAM^−^ cells were sorted by a FACSAria III cell sorter (BD Biosciences) after exclusion of CD90^+^, CD105^+^ and CD45^+^ cells.

#### RNA extraction and quantitative reverse transcription-PCR (qRT-PCR)

Total RNA was extracted from TGF-β1-treated cells (as described above) using NucleoSpin® RNA kit (Macherey-Nagel) according to manufacturer’s instructions.

Retrotranscription of RNA was performed using 3μg of total RNA in a 40μl final volume reaction mixture using iScript Advanced cDNA Synthesis Kit for RT-qPCR (Biorad) according to the manufacturer’s instructions. The reverse transcription reaction was processed at 42°C for 30min, and 85°C for 5min with a Mastercycler personal (Eppendorf).

qPCR was performed with StepOnePlus Real-Time PCR Systems (Applied Biosystems) in a reaction volume of 20μl containing 5μl of diluted cDNA, 2μM forward and reverse primers (Eurogentec), 1X MESA GREEN qPCR MasterMix Plus for Sybr® Assay (Eurogentec). qPCR was processed at 95°C for 5min, followed by 40 cycles of 95°C for 15s, 60°C for 30s and 72°C for 45s (data collection). All the qPCR reactions were performed in duplicate. The specificity of amplification was assessed for each sample by melting curve analysis, and the size of the amplicon was checked by electrophoresis of the PCR product on an agarose gel in TAE 1X (Life Technologies) (data not shown). All gene expression data were normalized with *hprt1* and *rplp0* expression data. Primer pairs were: 5’-CCCGAGTGTCAGCCTCAAA-3’ and 5’- GCAGGCTGCACACTGATCA-3’ for rat *Her2/Neu*, 5’-ATCCTCGCCCTGCTGATT-3’ and 5’- ACCACCGTTCTCCTCCGTA-3’ for *Cdh1*, 5’-GCGGCTCAGAGAGACTGTG-3’ and 5’- CCAAGCATTTAGACGCCAGTTT-3’ for *Epcam*, 5’-CCAACCTTTTCTTCCCTGAA-3’ and 5’- TGAGTGGGTGTCAACCAGAG-3’ for *Vim*, 5’-CTGACAGAGGCACCACTGAA-3’ and 5’- CATCTCCAGAGTCCAGCACA-3’ for *Acta2*, 5’-CGGTTTCACTTGAGAGCACA-3’ and 5’- CATACGTCCCAGGCTTTGAT-3’ for *Cdh2*, 5’-GCCAGCAGTCATGATGAAAA-3’ and 5’- TATCACAATACGGGCAGGTG-3’ for *Zeb1*, 5’-CCAGAGGAAACAAGGATTTCAG-3’ and 5’- AGGCCTGACATGTAGTCTTGTG-3’ for *Zeb2*, 5’-GTCTGCACGACCTGTGGAA-3’ and 5’- CAGGAGAATGGCTTCTCACC-3’ for *Snai1*, 5’-TTTCAACGCCTCCAAGAAGC-3’ and 5’- CGAGGTGAGGATCTCTGGTT-3’ for *Snai2*, 5’- CTGTGGACTGGCTCCATTTT-3’ and 5’- CAGTTTGATCCCAGCGTTTT-3’ for *Twist1*, 5’- AAACTGGACCAAGGCTCTCA-3’ and 5’- CAGGCTTCCTCGAAACAGTC-3’ for *Twist2*, 5’-TCCTCCTCAGACCGCTTTT-3’ and 5’- CCTGGTTCATCATCGCTAATC-3’ for *Hprt1*, 5’-ACTGGTCTAGGACCCGAGAAG-3’ and 5’- TCCCACCTTGTCTCCAGTCT-3’ for *Rplp0*.

#### Transcriptomic analyses of mouse mammary tumor cells

##### RNA-seq of tumor cells

Epithelial Her2^+^/EpCAM^+^ and mesenchymal Her2^+^/EpCAM^−^ tumor cells were FACS sorted from E-A14 tumors harvested from CD4/CD8 depleted mice and isotype control-injected (rat IgG2a/2b) mice, when tumors reached their endpoint volume (at day 20 and day 47 post tumor injection, respectively). Total RNA was extracted from sorted cells and from two replicated of *in vitro* cultured E-A14 cells stabilized in RLT buffer according to the “Purification of total RNA from animal and human cells” protocol of the RNeasy Micro Kit (Qiagen). The SMARTer Ultra Low Input RNA Kit for Sequencing v4 (Clontech Laboratories) was used to generate first strand cDNA from 0.1 to 1.0 ng total-RNA. Double stranded cDNA was amplified by LD PCR (10, 12 or 14 cycles) and purified via magnetic bead clean-up. Library preparation was carried out as described in the Illumina Nextera XT Sample Preparation Guide (Illumina). The libraries were quantified using the KAPA SYBR FAST ABI Prism Library Quantification Kit (Kapa Biosystems). Equimolar amounts of each library were pooled, and the pools were used for cluster generation on the cBot with the Illumina TruSeq SR Cluster Kit v3. The sequencing run was performed on a HiSeq 1000 instrument using the indexed, 50 cycles single-read protocol and the TruSeq SBS v3 Reagents according to the Illumina HiSeq 1000 System User Guide. Image analysis and base calling resulted in .bcl files, which were converted into .fastq files with the CASAVA1.8.2 software. RNA isolation, library preparation and RNAseq were performed at the Genomics Core Facility “KFB - Center of Excellence for Fluorescent Bioanalytics” (University of Regensburg, Regensburg, Germany). Sequencing reads were mapped to the *Mus musculus* genome sequence and transcriptome annotations (as provided by GENCODE release M9) and converted to gene counts using STAR (80). Sequencing reads mapping, transcriptome annotations and conversion to gene counts were performed by AltraBio (Lyon, France).

##### Statistical analyses of RNA-seq data

All statistical calculations were performed in R programming language (v. 3.5.3). DESeq2 (v. 1.22.2) (81) was used to normalize raw counts and to generate principal component analysis (PCA) plot based on the 500 most variable genes. The EMT score of each sample was determined by single sample gene set enrichment analysis (ssGSEA) of the EMT gene signature defined by Tan *et al*. (38) using GSVA (v. 1.30.0) (82). Differential expression analysis was performed using the DEseq2 package (v. 1.22.2) (81). Significantly differentially expressed genes (DEGs) between Her2^+^EpCAM^+^ and Her2^+^EpCAM^−^ tumor cells were identified with controlled False Positive Rate (B&H method) at 5% (False Discovery Rate (FDR) ≤ 0.05). Upregulated genes were selected at a minimum log2 fold change of 0.58 and downregulated genes at a minimum log2 fold change of −0,58. Hierarchical clustering of DEGs and samples was performed using the One-Pearson correlation as a metric and the complete linkage clustering method. The heatmap of DEG normalized expression values was performed using the Morpheus tool (Broad Institute). The clusterProfiler tool (v. 3.10.1) (83) was used for gene ontology (GO) and pathway analysis (KEGG, Reactome and Hallmark collections). Enriched gene sets were considered with a normalized enrichment scores (NES) > 1.5 and a FDR < 0.05.

##### Transcriptomic analyses of human breast tumors

The clinical outcome data and normalized expression dataset of breast invasive carcinoma (BRCA) from TCGA (33) (provisional data, February 2018) and from METABRIC (34) cohorts were downloaded from the cBioPortal (http://www.cbioportal.org/data_sets.jsp). For EMT signatures (EMT signature defined by Tan et al. (38) and Hallmark signature, https://www.gsea-msigdb.org/gsea/msigdb/collections.jsp), neutrophil signatures (signature from The human Protein Atlas, https://www.proteinatlas.org/, or signature defined by Ponzetta et al. (84)) and memory CD4 T cell signature (The human Protein Atlas), single sample enrichment scores were defined by ssGSEA analysis using the GSVA package. To evaluate the impact of CD4 T cells on tumor infiltration by neutrophils, human breast tumor samples were stratified into 3 groups (high, intermediate and low) based on tertiles of the memory CD4 T cell score. To evaluate the impact of tumor infiltration by neutrophils and CD4 T cells on EMT, human breast tumors samples were stratified to select 25% of samples with the highest (“high”) and 25% samples with the lowest (“low”) neutrophils and/or memory CD4 T cell scores. Graphical display of the spearman correlation matrix between all scores was performed using the corrplot R package.

##### Generation of cell culture conditioned mediums

E-A14, E-A21, 2G2 or M-A14 cell lines were cultured in DMEM supplemented with 5% FCS, 200Units.mL^−1^ penicillin, 200µg.mL^−1^ streptomycin and 1% L-glutamine. 48 hours after an appropriate cell seeding allowing to reach around 80% of cell confluence after two days, cell culture supernatants were harvested, centrifuged to remove remaining cells and cell debris and filtered with 0.2-micron syringe filter. Adherent cells were also counted and corresponding cell culture supernatant volumes were adjusted to take into account the slight cell growth differences between cell lines and to ensure an equal ratio between supernatant volume and adherent cell number across the different cell lines. These supernatants were then used for immune cell migration assay and cytokine quantification. For E-A14 and M-A14, two different supernatants of each (E-A14/SN#1 and E-A14/SN#2; M-A14/SN#1 and M-A14/SN#2) were produced during distinct cell culture experiments.

##### Immune cell migration assay

Immune cell migration was evaluated by a Transwell assay. Briefly, bone marrow cells were collected from tibiae and femora of FVB/N female mice at 8-10 weeks of age. After red blood cell lysis using BD Pharm Lyse™ (BD Biosciences), 5.10^5^ total cells were resuspended in 100µL DMEM supplemented with 5% FCS, 200Units.mL^−1^ penicillin, 200µg.mL^−1^ streptomycin and 1% L-glutamine, and loaded in each upper chamber of a transwell filter (polyester, 3µm pore diameter; Corning) inserted into a 24-well polystyrene plate. In the lower chambers, 600µL of the conditioned mediums from E-A14, E-A21, 2G2 or M-A14 cell lines was added. These supernatants were used from dilution 1:2 to 1:16. CXCL-1/KC/GROα was used as a chemoattractant positive control from 1µg.mL^−1^ to 250ng.mL^−1^. After 3 hours of incubation at 37°C under 5% CO_2_, the migrated cells were collected in the lower chambers and analyzed by flow cytometry using a CD45/CD11b/Ly6G/Ly6C/Gr1 multi-parametric staining. Absolute numbers of transmigrated cells were determined using Flow-count Fluorospheres (Beckman Coulter) according to the manufacturer’s instructions. Neutrophils were identified by their Ly6G^+^/Ly6C^int^ expression profile among CD45^+^ immune cells.

##### Cytokine quantification by MSD assay

CXCL1/KC/GROα was quantified in conditioned mediums using ECLIA (electrochemiluminescence assay) and MSD technology according to the U-plex protocol (MSD). Cell culture supernatants was harvested 48 hours after cell seeding.

##### Statistical analyses

To compare proportion of epithelial cells and neutrophils in tumors, statistical analyses were conducted using XLSTAT software. Differences between groups were analyzed with a Kruskal-Wallis *post-hoc* multiple pair wise comparison test using the *Steel*-*Dwass*-*Critchlow-Fligner* procedure.

To compare EMT score, statistical analyses were conducted using GraphPad Prism 6 with a non-parametric Wilcoxon matched pairs signed rank test.

*P* values less than 0.05 were considered significant (*, *P* < 0.05; **, *P* ≤ 0.01; ***, *P* ≤ 0.001; ****, *P* ≤ 0.0001).

## Data availability

The RNAseq datasets from murine tumor cell samples generated and analyzed in the current study are available in the GEO database under accession number GSE155577. METABRIC data are available in the EML-EBI archive (accession number: EGAS00000000083) and from supplementary information in Curtis *et al*. (34). TCGA BRCA RNAseq expression data were extracted as FPKM values from the GDC data portal (https://www.portal.gdc.cancer.gov/). All other data that supports the findings of this study are available within the article and its supplementary information files.

## ACKNOWLEDGEMENTS

The authors thank B.H. Nelson (University of British Columbia, Canada) for providing plasmid encoding the activated allele of rat *her2/neu*; K. De Visser (Netherlands Cancer Institute) for providing FVB RAG1KO mice; Genentech Inc. (California, U.S.A.) for providing anti-CD20 antibody; and M. Greene (University of Pennsylvania, U.S.A.) for providing 7.16.4 hybridoma. We thank AltraBio (Lyon, France) for the sequencing reads mapping of mouse tumor samples, transcriptome annotations and conversion to gene counts. We thank Manon Pratviel as well as the staff of the AniCan animal core facility for help in animal care and technical assistance. We also thank M. Ouzounova for proofreading the manuscript. This work was supported by funding from the Rhône and Ardèche comity of the “Ligue Contre le Cancer”, the Fondation ARC pour la Recherche sur le Cancer, the Institut National contre le Cancer (project PLBIO15-266) and the SIRIC LYriCAN project (grant INCa-DGOS-Inserm_12563). This work was additionally supported by funding from the LABEX DEVweCAN (ANR10-LABX-0061) of Université de Lyon, within the program “Investissements d’Avenir” (ANR-11-IDEX-0007) operated by the French National Research Agency (ANR) and the Institut Convergence Plascan (Grant Number ANR-17-CONV-0002). We would like to thank our financial supports: the “Ligue Nationale Contre le Cancer” for A.S. and the Auvergne-Rhône-Alpes region for S.A.

## AUTHOR CONTRIBUTION

A.S., C.C. and I.P. designed the research project. A.S., D.P., S.A., C.P., N.G., M.M. and M.C.C. performed experiments. A.S., S.A., M.H., C.P., J.P.F., M.C.C., C.C. and I.P. analyzed data; I.D. performed cell sorting by flow cytometry; P.S. provided helpful discussions; A.P. provided strategic advice, critical evaluation of the manuscript and scientific content. A.S., M.H., C.C. and I.P. wrote the manuscript. C.C. and I.P. supervised the research. All authors read and approved the final version of the manuscript.

## COMPETING INTERESTS

The authors declare no potential conflicts of interest.

## Notes

### Competing Interest Statement

The authors have declared no competing interest.

