## supplemental figures and legends for "CD4 T cells and neutrophils contribute to epithelial-mesenchymal transition in breast cancer"

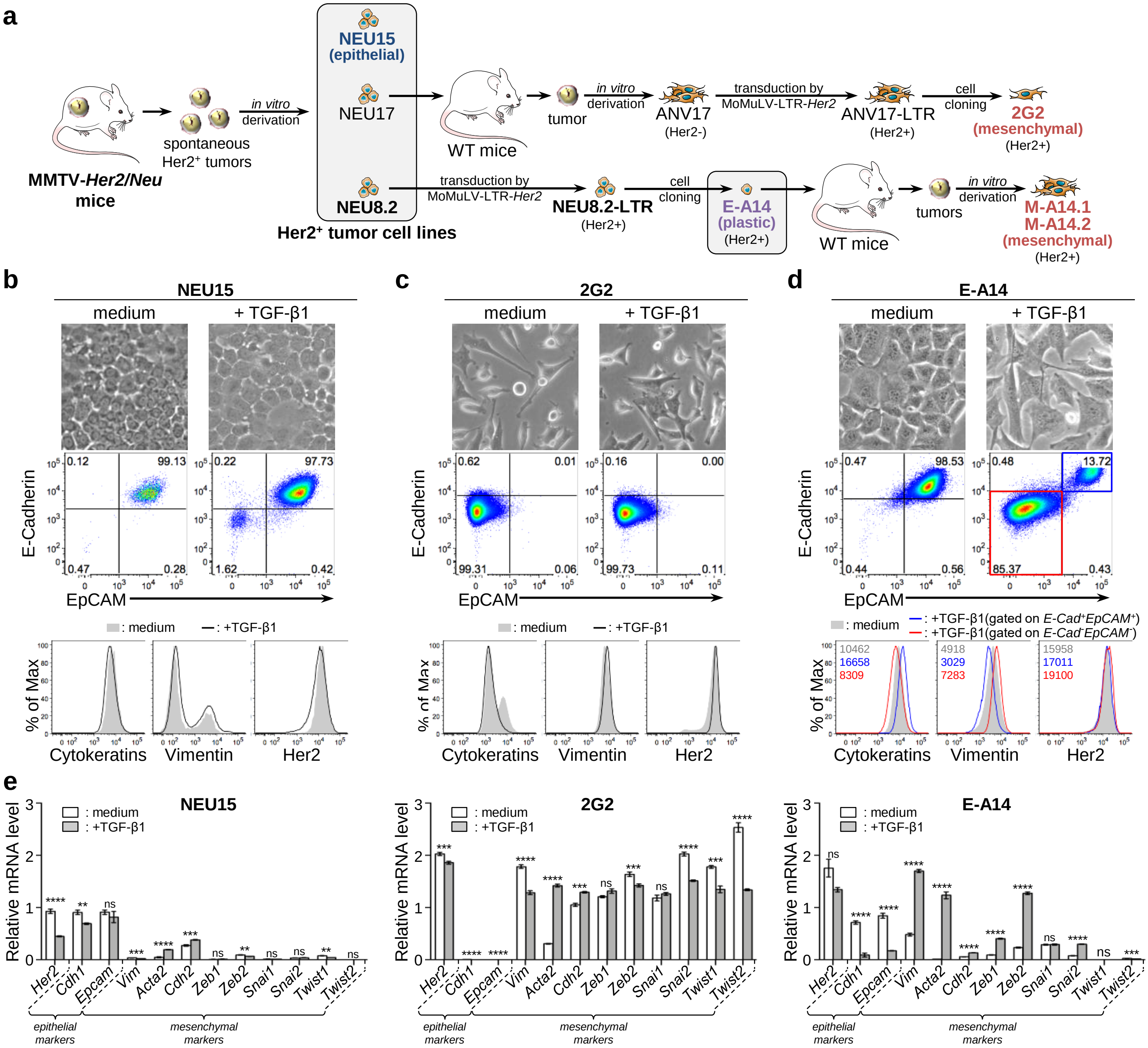


| | NEU15 | 2G2 | E-A14 |
|---|---|---|---|
| *Her2* | 0.49 ± 0.02 | 0.92 ± 0.02 | 0.77 ± 0.07 |
| *Cdh1* | 0.77 ± 0.03 | 0.09 ± 0.01 | 0.12 ± 0.05 |
| *Epcam* | 0.90 ± 0.11 | 2.80 ± 0.20 | 0.20 ± 0.01 |
| *Vim* | 0.59 ± 0.03 | 0.72 ± 0.02 | 3.52 ± 0.19 |
| *Acta2* | 4.08 ± 0.87 | 4.60 ± 0.14 | 81.25 ± 4.21 |
| *Cdh2* | 1.40 ± 0.07 | 1.23 ± 0.03 | 2.30 ± 0.06 |
| *Zeb1* | 1.58 ± 0.24 | 1.09 ± 0.04 | 4.24 ± 0.21 |
| *Zeb2* | 0.73 ± 0.05 | 0.87 ± 0.02 | 5.39 ± 0.25 |
| *Snai1* | 0.92 ± 0.05 | 1.07 ± 0.05 | 1.00 ± 0.06 |
| *Snai2* | 1.07 ± 0.05 | 0.75 ± 0.01 | 3.76 ± 0.18 |
| *Twist1* | 0.55 ± 0.03 | 0.76 ± 0.03 | 2.71 ± 0.60 |
| *Twist2* | 0.76 ± 0.14 | 0.53 ± 0.02 | 0.61 ± 0.06 |

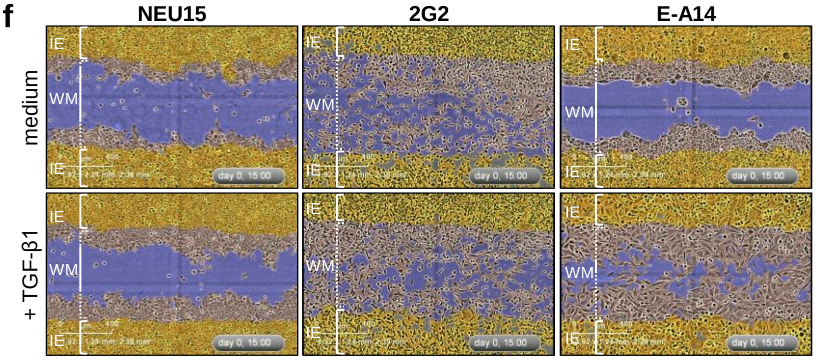


**Supplementary Fig. 1: Comparative phenotype of epithelial, mesenchymal and plastic mammary tumor cell lines**

**a-** Generation of Her2-expressing mouse mammary tumor cell lines. **b-f-** NEU15, 2G2 and E-A14 mammary tumor cells were treated *in vitro* with TGF-β1. **b-d-** The morphology of cells was visualized using light microscopy (original magnification x200). Cells were also analyzed by flow cytometry for the expression of E-Cadherin, EpCAM, Cytokeratins, Vimentin and Her2. Gates are based on the isotype control corresponding to each marker. Numbers on histograms indicate mean fluorescence intensity. **e-** *Her2*, *Cdh1*, *Epcam*, *Vim*, *Acta2*, *Cdh2*, *Zeb1*, *Zeb2*, *Snai1*, *Snai2*, *Twist1* and *Twist2* mRNA expression was quantified by RT-qPCR. Graphs show gene expression normalized by *Hprt1* and *Rplp0* housekeeping genes. Bars and error bars represent the mean and standard deviation, respectively. n=4. Student's t test. **, ***, ****: p ≤ 0.01, 0.001 or 0.0001. Table data indicates gene expression fold change (mean ± standard deviation) in TGF-β1-treated cells in comparison with control (medium) cells, with a color scale running from minimal (blue) to maximal fold change value (red). **f-** Cells were grown in a monolayer in TGF-β1-supplemented medium. A scratch wound was made and images were taken using the Incucyte ZOOM instrument (Essen BioScience). Pictures presented here display wound healing after 15h. The blue region denotes the scratch wound mask (WM) over time as cells migrate into the wound region. The initial migrating/invading edges (IE) of the wound, created immediately following wound creation, are shown in yellow.

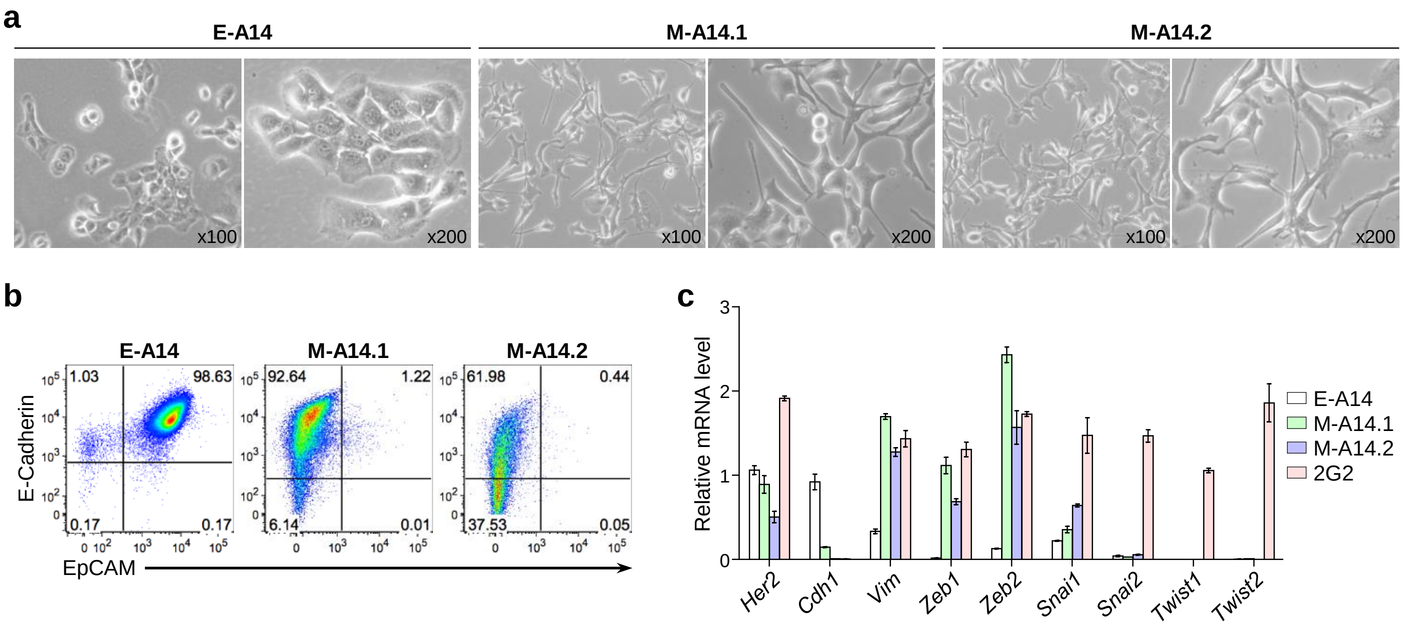


**Supplementary Fig. 2: Tumors that emerge in adaptive immunocompetent mice display mesenchymal features**

**a-** Bright field images of E-A14 and two different cell lines, M-A14.1 and M-A14.2, derived from tumors arisen in WT mice after E-A14 injection. Cells were also analyzed by **b-** flow cytometry for E-Cadherin and EpCAM expression and by **c-** quantitative RT-PCR for the mRNA expression of epithelial and mesenchymal genes. **b-** Gates are based on the isotype control corresponding to each marker. **c-** Graph shows gene expression normalized by *Hprt1* and *Rplp0* housekeeping genes. Bars and error bars represent the mean and standard deviation, respectively. n=2.

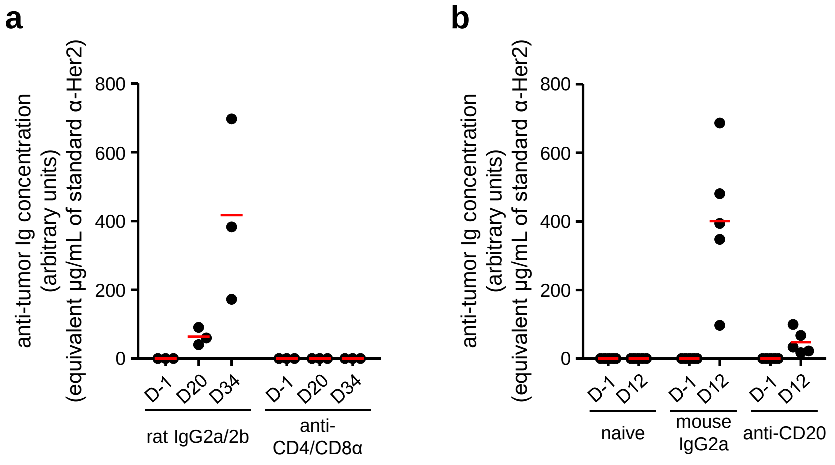


**Supplementary Fig. 3: Humoral response against E-A14 tumor cells**

E-A14 mammary tumor cells were injected into the mammary fat pad of **a-** CD4/CD8α double-depleted or isotype (rat IgG2a and IgG2b) control-injected mice, and **b-** CD20-depleted mice or isotype (mouse IgG2a) control-injected mice. Specific anti-E-A14 tumor immunoglobulin (Ig) level was measured, in the plasma of E-A14 tumor-bearing mice, the day prior tumor cell injection (D-1) and at different time points during tumor growth: **a-** 20 (D20) and 34 (D34) days after tumor injection; **b-** 12 (D12) days after tumor injection. **b-** Ig concentration was also measured in serum samples harvested at the same time in non-tumor bearing (naive) mice. Concentration of anti-tumor antibodies was calculated using a standard curve constructed with a standard anti-Her2 antibody. Red bars on the graph indicate the mean for each group. **a-** n=3 mice for each group. **b-** n=5 mice for each group.

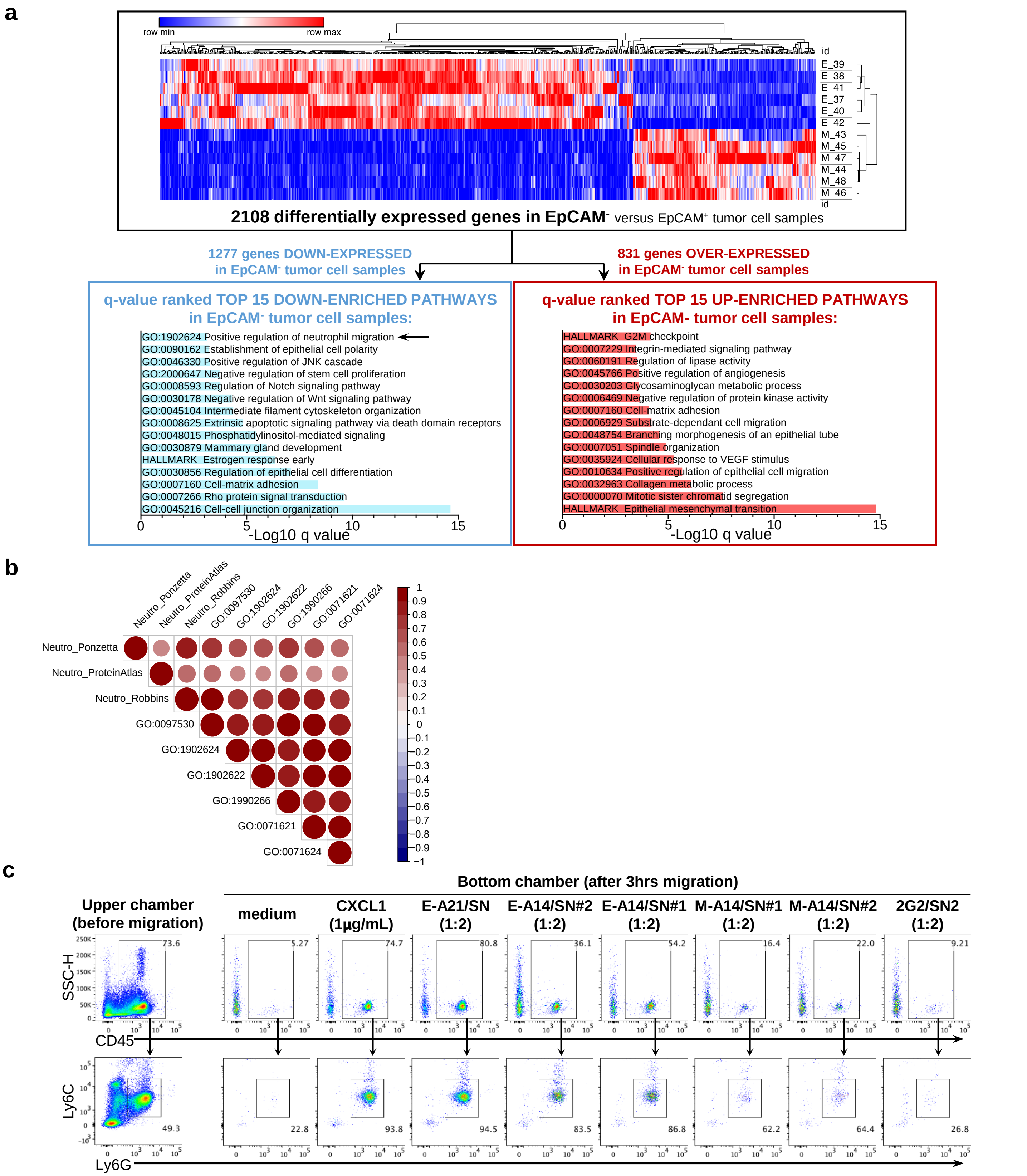


**Supplementary Fig. 4: Differential neutrophil recruitment ability between epithelial and mesenchymal tumor cells**

**a-**Transcriptomic profile of FACS-sorted EpCAM+ (E) and EpCAM- (M) tumor cells, isolated from E-A14 tumors harvested in isotype (rat IgG2an and IgG2b) control-injected mice (n=6), at tumor end-point (47 days after tumor cell injection). Top panel: hierarchical clustering of all significantly differentially expressed genes (DEGs) (n=2108 DEGs) between EpCAM+ (E) and EpCAM- (M) tumor cell samples. DEGs analysis identified 831 genes significantly over-expressed and 1277 genes down-expressed in EpCAM- compared to EpCAM+ tumor cells. Bottom panel: pathway enrichment analysis of DEGs between EpCAM+ and EpCAM- tumor cell samples. The top 15 most down- (bottom left panel) and up- (bottom right panel) enriched gene pathways in EpCAM- (versus EpCAM+) tumor cells are presented here. **b-** Correlogram based on the Spearman rank correlation between the different scores of neutrophil signatures and GO terms related to neutrophil or granulocyte chemotaxis/migration, determined by ssGSEA for human tumors samples from METABRIC cohort. **c-** Immune cell chemotaxis induced by epithelial (E-A21, E-A14) or mesenchymal (M-A14, 2G2) tumor cell line supernatants used in Fig. 5e-f. CXCL1 was used as a positive control. For E-A14 or M-A14, two different supernatants (SN#1 and SN#2) were produced during distinct cell culture experiments. Ly6G+Ly6C$^{int}$ neutrophil frequency (bottom panel) was determined among total migrating CD45+ immune cells (top panel). Gates are based on the isotype control corresponding to each marker. A representative dot plot for each condition is presented, n=3. Data are representative of two independent experiments.

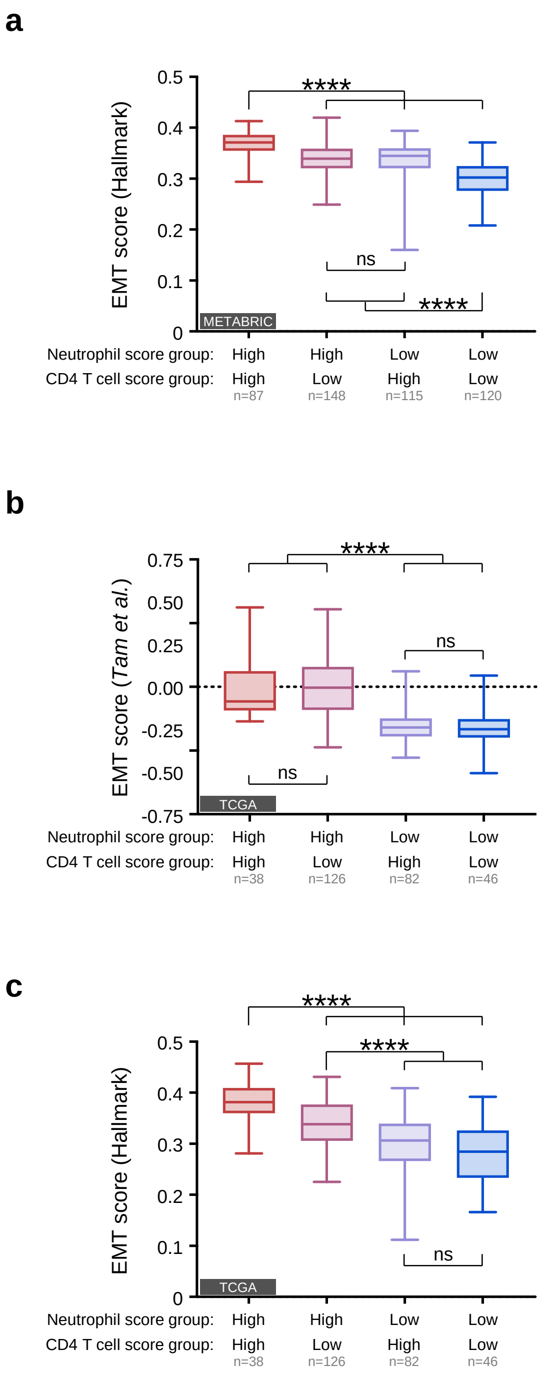


**Supplementary Fig. 5: Single sample enrichment scores of EMT signatures in transcriptomic data of breast tumors from METABRIC and TCGA cohorts.**
Tumors from **a-** METABRIC or **b-c-** TCGA cohorts were selected according to their neutrophil and memory CD4 T cell enrichment scores (signatures both from The human Protein Atlas), both determined by ssGSEA, with high and low groups defined by the 25% of tumors with the highest and the lowest scores. Enrichment score for **a, c-** EMT Hallmark signature or **b-** EMT signature defined by *Tan et al., EMBO Molecular Medicine 2014*, were also determined by ssGSEA. Kruskal-Wallis test. *, ****: p ≤ 0.05 or 0.0001.

**Supplementary Files 1-6. Time-lapse videos of migration assay using NEU15, 2G2 and E-A14 mammary tumor cells**
NEU15, 2G2 and E-A14 cells were grown in a monolayer in medium, supplemented or not with TGF-β1. A scratch wound was made and the scratch area was monitored using the Incucyte Live-cell Imaging System (Essen Bioscience). Images were acquired every 30min using a 10x objective and videos were exported using Incucyte ZOOM software. The blue region denotes the scratch wound mask over time as cells migrate into the wound region. The initial migrating/invading edges of the wound, created immediately following wound creation, are shown in yellow.

**Supplementary File 1. Time-lapse video of migration assay using NEU15 cells in control condition.**

**Supplementary File 2. Time-lapse video of migration assay using NEU15 cells in TGFβ1-treated condition.**

**Supplementary File 3. Time-lapse video of migration assay using 2G2 cells in control condition.**

**Supplementary File 4. Time-lapse video of migration assay using 2G2 cells in TGFβ1-treated condition.**

**Supplementary File 5. Time-lapse video of migration assay using E-A14 cells in control condition.**

**Supplementary File 6. Time-lapse video of migration assay using E-A14 cells in TGFβ1-treated condition.**
